# The effect of habitat loss and fragmentation on isolation-by-distance and time

**DOI:** 10.1101/2022.10.26.513874

**Authors:** Gabriele Maria Sgarlata, Tiago Maié, Tiago de Zoeten, Rita Rasteiro, Lounès Chikhi

## Abstract

Throughout Earth’s natural history, habitats have undergone drastic changes in quality and extent, influencing the distribution of species and their diversity. In the last few hundred years, human activities have destroyed natural habitats at an unprecedent rate, converting continuous habitat into fragmented and isolated patches. Recent global metanalyses suggest that habitat loss and fragmentation (HL&F) has negatively impacted the genetic diversity of many taxa across the world. These conclusions have been drawn by comparing present-day genetic patterns from populations occurring in continuous and fragmented landscapes. In this work, we attempted to go beyond ‘pattern’ and investigate through simulations some of the ‘processes’ that influence genetic variation in the context of HL&F. Since most species have a geographically restricted dispersal (known as “isolation-by-distance”, IBD), we studied the impact of HL&F on isolation-by-distance. We characterised the behaviour of IBD in the case of i) instantaneous HL&F, ii) gradual (two-steps) HL&F, and iii) instantaneous HL&F following range expansion. In addition, we propose a spatially-explicit theoretical framework by modifying the original theoretical results on isolation-by-distance (Slatkin, 1991; Slatkin, 1993) and apply them to a toroidal stepping-stone model in the context of HL&F. Our results suggest that isolation-by-distance can be maintained for relatively long time after HL&F, thus pointing to the long-term importance of spatial genetic structure in species genetic diversity. In addition, our results may explain why present-day fragmented population still show significant IBD pattern although being disconnected.

## Introduction

Throughout Earth’s natural history, environmental changes have influenced the distribution and abundance of species worldwide, for instance, through habitat contraction and expansion (Hewitt, 2000; Newbold et al., 2015; Ceballos et al., 2017; Cooper et al., 2021). In the last two centuries, human activities have intensified the destruction and degradation of natural ecosystems, often leading to habitat loss and fragmentation (HL&F), a process characterized by a loss in the amount of habitat area (*habitat loss*) and the splitting of habitat into smaller isolated patches (*habitat fragmentation per se*) (Curtis, 1956; Moore, 1962; Fahrig, 2003).

Multiple approaches allow today to investigate species response to HL&F, for example i) by reconstructing the natural history of species and their environment (Hewitt, 1996; Barak et al., 2016; Lyman, 2017; Grace et al., 2019), ii) by comparing populations occurring in landscapes with different degree of habitat disturbance (e.g., Lino et al., 2019; Almeida-Rocha et al., 2020) or iii) by performing long-term experiments (e.g., Haddad et al., 2015, 2017). Genetic data are one of the sources of information that are used to reconstruct the natural history of species (i.e., past changes in population size and connectivity) (Nielsen and Beaumont, 2009; Beichman et al., 2018; Mitchell and Rawlence, 2021; Loog, 2021) and, together with data on the history of habitat changes (e.g., paleoenvironmental data), can provide crucial insights on how species have responded to past HL&F (e.g., Teixeira et al., 2021). Consequently, learning about past species response to HL&F, can contribute to build predictions on how species will be affected by present-day HLF.

However, the history that genetic data can recover relies on the assumptions of the models that are used for demographic inference. In particular, there are frameworks assuming that species are not structured (e.g., Beaumont, 1999; Li and Durbin, 2011; Nikolic and Chevalet, 2014; Liu and Fu, 2015), namely that individuals reproduce at random and that mating can occur with equal probability with any of the individuals in the population (“*panmictic* population”). Other methods consider that species might be subdivided in smaller panmictic subpopulations (‘islands’ or ‘demes’), all connected with each other by gene flow (“*structured* population”) (e.g., Nielsen and Wakeley, 2001; Hey and Nielsen, 2004; Gutenkunst et al., 2009; Costa and Wilkinson-Herbots, 2017; Arredondo et al., 2021). However, relatively few inferential approaches take explicitly into account the form of population structure that is probably most common in the wild, that is *spatial population structure* (Meirmans, 2012; Sexton et al., 2014; Aguillon et al., 2017). This form of spatially-explicit population structure considers that dispersal (and thus gene-flow) is more likely with close-by subpopulations than with those far apart.

Geographically-limited dispersal generates a spatial genetic pattern called ‘isolation-by-distance’ (IBD) (Wright, 1943; Malécot, 1948). Since the seminal works of Wright, (1940, 1943, 1946) and Malécot, (1948), several spatial models have been proposed to explain spatial genetic variation over short (Barton and Wilson, 1995; Wilkinson-Herbots, 1998; Barton et al., 2002; Cox and Durrett, 2002) and large geographic scales (Barton et al., 2010). Nevertheless, spatial genetic models typically assume continuous (i.e., not fragmented) habitat and, homogenous population density and dispersal. Some exceptions are, for instance, Robledo-Arnuncio and Rousset, (2010), which derived exact recursion equations for IBD model allowing variation in local population density, and Barton, (2008), which constructed a theoretical framework describing the effect of a physical barrier on isolation-by-distance pattern. More recent studies have open new avenues of research in spatial population genetics, going beyond some of the assumptions initially formalized in the classical ‘isolation-by-distance’ model (e.g., homogenous dispersa; McRae, 2006), and recovering information on genetic ancestry at finer geographical- and temporal-scale (Ralph and Coop, 2013; Ringbauer et al., 2017). For instance, Duforet-Frebourg and Slatkin, (2016) have introduced the theory of isolation-by-distance and time (IBDT), a generalization of the IBD theory, by allowing differences in sampling time of the genetic data. Nevertheless, to date, there are no spatially-explicit frameworks that allow to consider the joint effect of geographically-limited dispersal (space) and divergence (time) on genetic diversity.Filling this knowledge gap may indeed be important for better understanding how HL&F influences the genetic diversity of species, since HL&F is a spatial-temporal process (Mandelbrot, 1982; Krummel et al., 1987; Milne, 1988; Solé and Manrubia, 1995) and given that most species are characterised by geographically-limited dispersal.

In the present study, we investigated non-equilibrium IBD patterns in a two-dimensional population (*landscape*) undergoing HL&F. According to Hutchison and Templeton (1999), HL&F would result into a lack of correlation between genetic and geographical distance, due to the stronger influence of local drift compared to gene-flow among fragmented populations (case III in Hutchins and Templeton, 1999). However, several genetic studies have shown that significant IBD signatures can still be detected among fragmented subpopulations, although violating the assumption of continuous habitat in the IBD model. Some examples are the northern rufous mouse lemur (Aleixo-Pais et al., 2019), the marsh fritillary (Pertoldi et al., 2021), the white carob tree (Roser et al., 2017), the Asian rice (Zhao et al., 2013), the blue duck (Grosser et al., 2017) and the long-lived subalpine conifer (Tóth et al., 2019). Assuming that there is no ongoing gene-flow among fragmented patches, these studies may suggest that genetic data keep ‘memory’ of a past in which the landscape was continuous and not-fragmented. But for how long this ‘memory’ can be maintained? Or more in general, how IBD patterns change after HL&F?

In this study we used a simulation-based approach to explore the behaviour of IBD over time under different scenarios of HL&F. More specifically, we ask *i)* what is the spatio-temporal dynamic of IBD in a two-dimensional landscape undergoing *instantaneous HL&F*, and *ii)* how range expansion and variation in tempo and mode of HL&F influences the persistence of IBD patterns. We conclude proposing a theoretical framework that consider both the effect of space and time in shaping population genetic variation in the context of HL&F. In particular, we modify the original theorical results on isolation-by-distance (Slatkin, 1991, 1993) and apply them to a toroidal stepping-stone model undergoing HL&F. Our analytical model opens the possibility for future development of an inferential-approach allowing to interpret *F*_*st*_ pattern considering both space and time.

## Materials and Methods

### Population genetics simulations

We used the individual-based computer program SINS (Simulating INdividuals In Space; Rasteiro et al., 2012) to model a two-dimensional landscape experiencing habitat loss and fragmentation. SINS uses a forward-time approach to simulate diploid individuals (males and females) over a 2D grid of demes, similar to a classical 2D stepping-stone model (Kimura and Weiss, 1964) and inspired by the original coalescent-based software SPLATCHE (Currat et al., 2004). SINS and SPLATCHE share several features, but they are also complementary since they differ in some properties. More details on the comparison between SINS and SPLATCHE can be found in Rasteiro et al., (2012).

In SINS, deme population size 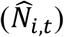 is drawn from a Poisson distribution with a mean value (*N*_*i,t*_) given by a corrected form of the Maynard-Smith and Slatkin, (1973) logistic growth. The *N*_*i,t*_ value in each deme is logistically regulated by deme-specific carrying capacity (*K*), growth rate (*r*) and deme number of reproductive females (*N*_*f*_). Males and females mate at random within demes. Each deme exchanges migrants with the four neighbouring demes. The number of individuals that will emigrate from the focal deme at each generation is drawn from a Poisson distribution with mean *M* = *N*_*i,t*_*m n*_*d*_/4, where *m* is the dispersal rate and *n*_*d*_ is the number of available neighbouring demes (e.g., 2 for a corner deme and 4 for central deme). The number of migrants will then be stochastically distributed among the neighbouring demes using a binomial distribution with probability *P*_*dir*_ = (1 − *F*_*dir*_)/(*n*_*d*_ − *F*_*t*_), where *dir* is one of the available directions (min=1; max=4), *F*_*dir*_ is a deme-specific friction parameter defining the difficulty to move from the focal deme into that direction and *F*_*t*_ is the sum of the frictions of the *n*_*d*_ receiving demes. Similar to SPLATCHE, changes from one landscape configuration to another (i.e., continuous to fragmented) are defined by *K* and *F* maps describing, respectively, the carrying capacity and friction value of each deme in the two configurations. Demes that are unsuitable after an ‘environmental event’ are set with *K* = 0 and *F* = −99, such that no individuals can grow and no migrants can move within these unsuitable demes. The growth rate was set to 0.8, meaning that fraction of the carrying capacity available for further growth will generate 0.8 offspring per generation.

SINS is able to simultaneously simulate demographic and genetic data over non-overlapping discrete generations and through space. In the present study we will exclusively use independent microsatellite loci, which mutations are modelled according to the Stepwise Mutation Model (Ohta and Kimura, 1973). Simulations start with all demes having the same distribution of allele frequencies. We set initial allele frequency for each locus by sampling 100 individuals from a Wright-Fisher (WF) population simulated in *fastsimcoal* (Excoffier and Foll, 2011) using the *fastsimcoal* function in the *strataG* R package (Archer et al., 2017). The size of the WF population was set equal to 10,000 and mutation rate according to the value specified in the SINS simulation. We decided to define the initial deme allele frequencies in order to ensure that dispersal-mutation-drift equilibrium would be reached quicker than if we had to wait for mutations to occur. Three scenarios were simulated in this study:

#### Instantaneous HL&F

We simulated a two-dimensional grid of 13 × 13 demes (*d*_*tot*_ = 169 demes), with homogenous deme-population size (*K*) and deme-specific friction (*F*). After 10,000 generations, more than the time required for reaching a ‘quasi-genetic equilibrium’ (i.e., Δ*He*/*t* ≈ 0), the landscape is subjected to HL&F, leading to nine isolated patches of 3 × 3 demes (Fig. 1a). After that, simulations were run for over 4,000 generations. We tested several values of *K* (50, 100, 200) and *m* (0.02, 0.04, 0.06, 0.1), which were kept constant across all demes in the landscape. We also tested several values of mutation rate, *μ* (5 × 10^−3^, 5 × 10^−4^, 5 × 10^−5^, 5 × 10^−6^) and number of genetic markers (15, 30, 100, 400 microsatellites).

**Figure 1.**
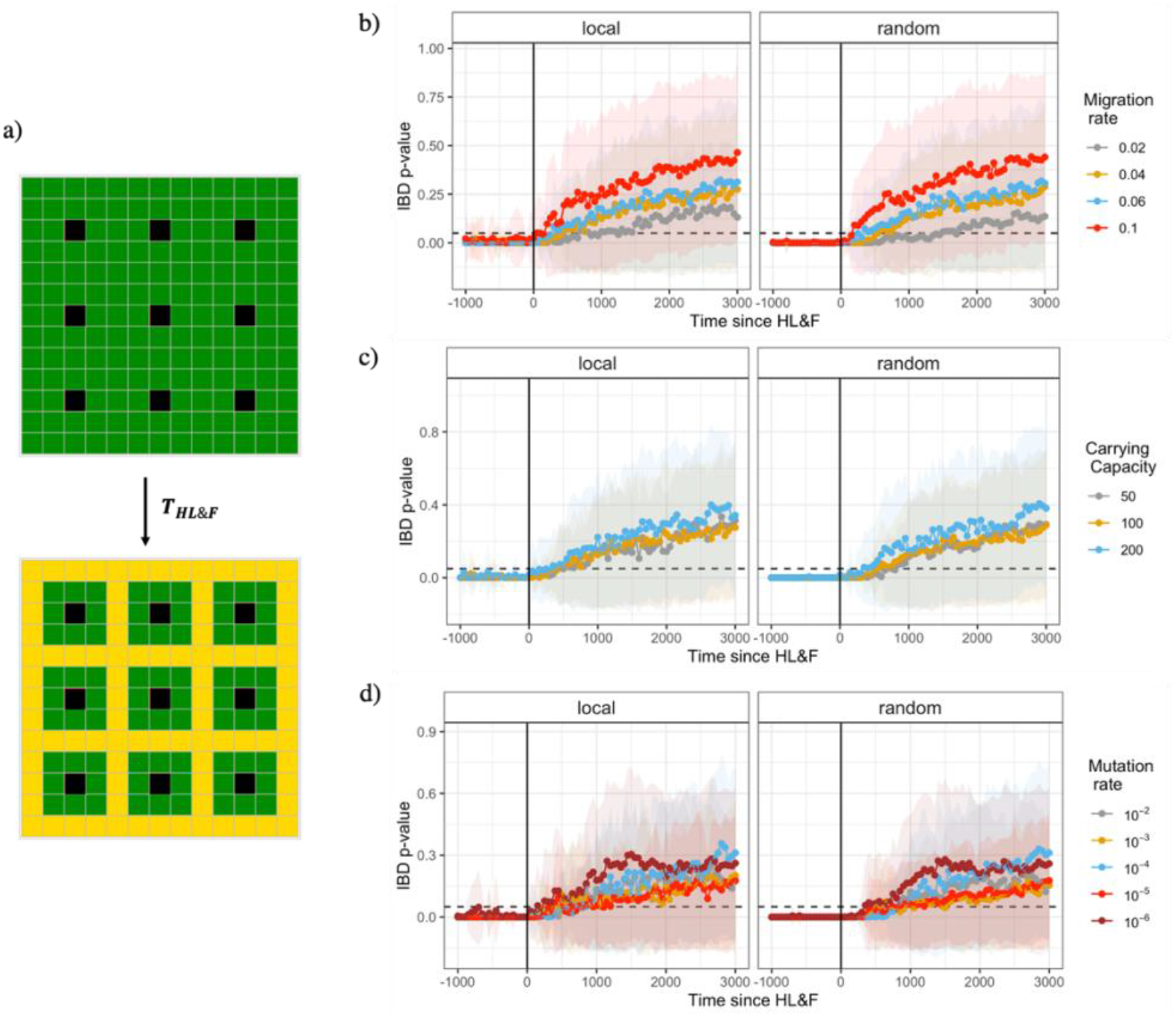
The effect of habitat loss and fragmentation on *R*_*st*_-based isolation-by-distance (IBD) in relation to dispersal rate, population size and mutation rate. **a)** Simulated 13 × 13 square grid of demes (*d*_*tot*_ = 169) undergoing HL&F at time *T*_*HL&F*_. Black cells refer to the local sampling scheme, green and yellow cells refer to suitable and unsuitable habitat, respectively. Time at which IBD is no longer significant (*T*_*IBD*_) as a function of **b)** dispersal rate (*m* = 0.02− 0.01, *K* = 100, *μ* = 5 × 10^−4^), **c)** population size (*K* = 50 − 200, *m* = 0.04, *μ* = 5 × 10^−4^) and mutation rate (*μ* =10^−2^-10^−6^, *K* = 100, *m* = 0.02). Mean (dots) and variance (shaded area) of IBD p-value across simulation replicates. Dashed line represents p-value = 0.05. Local sampling: patch-sampled individuals come from the same deme; Random sampling: patch-sampled individuals are randomly collected across demes within the patch. The results suggest i) that IBD is lost quicker at high than low dispersal rate, ii) that population size has little effect on *T*_*IBD*_, and iii) that mutation rate does not influence *T*_*IBD*_. Note, however, a large variance in *T*_*IBD*_ across all simulated parameter combinations.

#### Range expansion prior to instantaneous HL&F

Range expansion is an evolutionary process that has shaped the genomic diversity of many taxa across the world (Hewitt, 2000; Eckert et al., 2008; Pauls et al., 2013). It is thus relevant to understand how prior range expansion genetic signatures influence genetic pattern generated by HL&F. The simulated scenario is equivalent to the *instantaneous HL&F*, apart from the initial conditions. In fact, the 13 × 13 grid is initially unoccupied by individuals, except in the lower-left corner deme. From such deme, individuals are sent to the neighbouring demes according to the simulated dispersal rate (*m*). We assessed the impact of past range expansion before the instantaneous HL&F event by setting the time of HL&F at different time points since the onset of the range expansion (the starting of the simulation; Δ*t*_*exp*_): 750, 1000 and 1500 generations (Fig. 2a). The minimum Δ*t*_*exp*_ value (= 750) was chosen such to ensure complete colonisation of the 13 × 13 landscape before HL&F.

**Figure 2.**
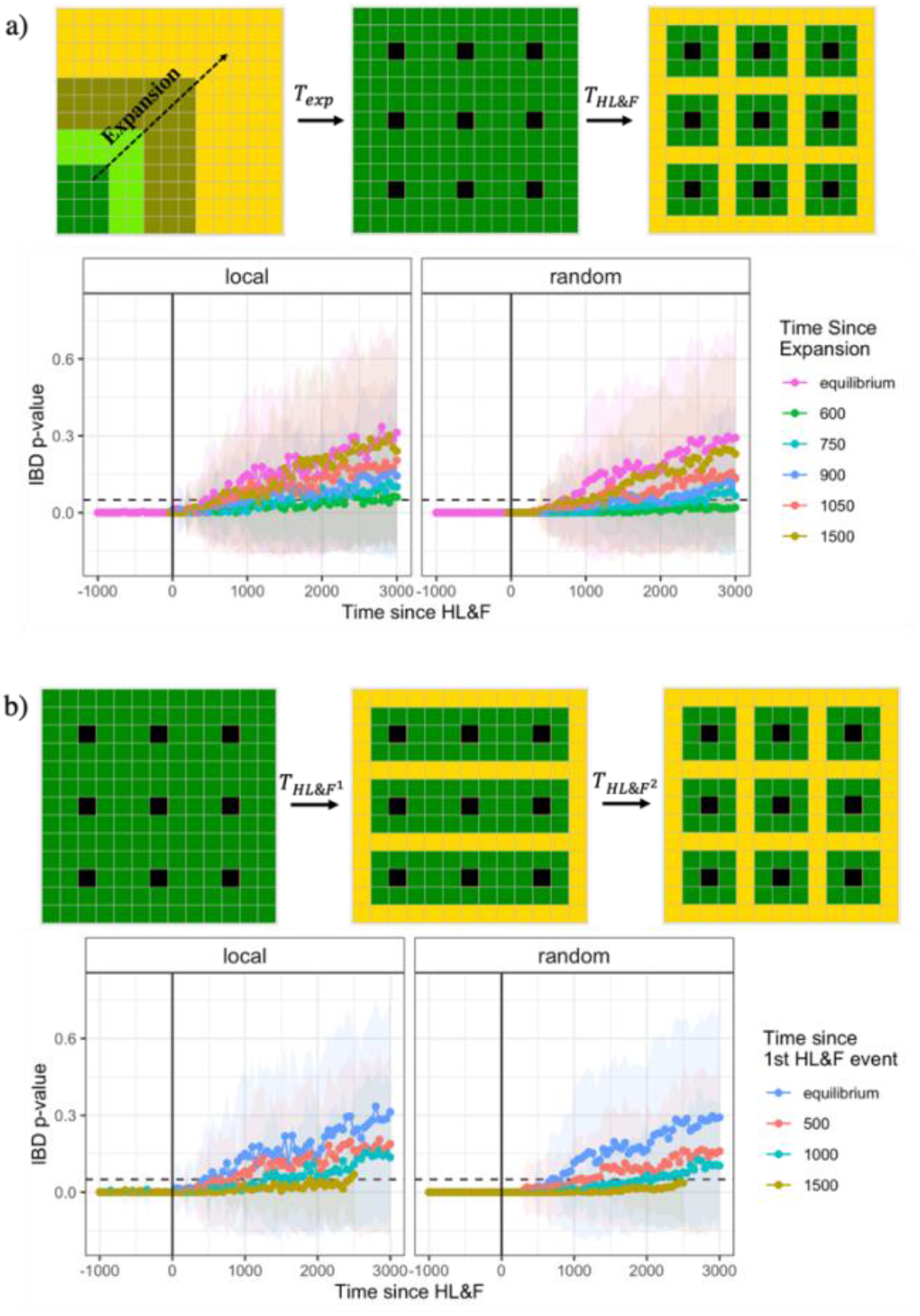
The effect of ancient range expansion followed by HL&F or spatio-temporal variation in HL&F on loss of R_**st**_-based isolation-by-distance. **a)** Simulated 13 × 13 square grid of demes, where *T*_*exp*_ is the time since the onset of the range expansion and *T*_*HL&F*_ is the time of instantaneous HL&F. Black cells refer to the local sampling scheme, green and yellow cells refer to suitable and unsuitable habitat, respectively. Range expansion was simulated for a scenario with *K* = 50, *m* = 0.04 and *μ* = 5 × 10^−4^. The results show that the genetic drift originated during a recent range expansion can contribute to a longer persistence of IBD following a HL&F event. **b)** Simulated 13 × 13 square grid of demes, where 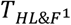 and 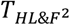 indicate, respectively, the time of the 1^st^ and 2^nd^ HL&F event. Piecewise HL&F was simulated for a scenario with *K* = 50, *m* = 0.04 and *μ* = 5 × 10^−4^. The results show that spatio-temporal variation in HL&F can have a strong effect on *T*_*IBD*_, which is exacerbated with increasing Δ*T*_*HL&F*_. Mean (dots) and variance (shaded area) of IBD p-value across simulation replicates. Dashed line represents p-value = 0.05. Local sampling: patch-sampled individuals come from the same deme; Random sampling: patch-sampled individuals are randomly collected across demes within the patch.

#### Two-steps HL&F

In real landscape, HL&F may not necessarily occur instantaneously. Instead, it is more likely that habitat fragments are formed gradually following fine-scale spatial-temporal processes (Mandelbrot, 1982; Solé and Manrubia, 1995). Therefore, we assessed the impact of gradual HL&F by simulating a scenario where HL&F occurs as a two-steps event. The initial two-dimensional landscape is equivalent to the initial conditions of the *Instantaneous HL&F* scenario, before HL&F. After 10,000 generations, the landscape is subject to a first HL&F event 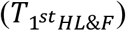, forming three large isolated habitat fragments that undergo a second HL&F event after Δ*t*_2*n*__*d*_ generations. The second HL&F event produces a fragmented landscape with nine isolated patches of 3 × 3 demes, equivalent to the *instantaneous HL&F* scenario (Fig. 2b). We considered three values of Δ*t*_2*n*__*d*_: 500, 1000 and 1500 generations.

Across all simulations, we sampled 14 individuals at 30 microsatellites from each isolated patch (*N*_*tot*_: 14 × 9 = 126 individuals), before and after HL&F. Two sampling strategies are considered: *local sampling*, where individuals are exclusively sampled at the central deme of each patch; and *random sampling*, where individuals are randomly sampled across the 9 demes of each isolated patch. We ran 100 independent simulation replicates for each scenario and recorded genetic data every 50 generations.

### Measuring spatial genetic patterns

Pairwise genetic differentiation was estimated using the *R*_*st*_ statistic, since it is more appropriate for microsatellite data mutating according to the Stepwise Mutation Model (Slatkin, 1995). We computed *R*_*st*_ using the *polysat* R package (Clark and Jasieniuk, 2011). Genetic differentiation was estimated for each independent microsatellite locus and then averaged across loci. Isolation-by-distance (IBD) was computed by correlating pairwise *R*_*st*_ /(1 − *R*_*st*_) with the log-transformed geographic distance as proposed in Rousset, (1997) for two-dimensional habitats. Pairwise geographic distances were calculated using the *euclidean* distance. The significance of IBD was assessed using Mantel test. For comparison, we also estimated genetic differentiation using Wright’s *F*_*st*_.

The quantification of the time of IBD loss (*T*_*IBD*_) was carried out on the time series IBD p-value for each simulation replicate by using a changepoint method based on PELT-algorithm, implemented in the *changepoint* R package (Killick and Eckley, 2014). It estimates the point at which the statistical properties of a (temporal) series of observations (p-value in this case) change. We used the *cpt*.*mean* function (Killick et al., 2011) to identify a single change point of the mean signal over time. The function segments the data in intervals of a user-defined size (penalty value) and, for each interval, computes the average data value. Different penalty values (1 – 10) are iteratively tested until a detection point (*T*_*IBD*_) is found.

Relative Weights Analysis (RWA) was used to quantify the relative contribution of geographical distance and time since HL&F on pairwise *R*_*st*_ (or *F*_*st*_). RWA is a statistical method that computes the relative importance of predictor variables on the response variable of a regression model, accounting for possible correlations among predictor variables (Tonidandel et al., 2009). RWA was carried out using the *rwa* R package (Chan, 2022).

### A toroidal stepping-stone model under HL&F

Isolation-by-distance can be visualized by linear regression of deme pairwise *F*_*st*_ (or *R*_*st*_ if using microsatellites) and geographical distance. Alternatively, but formally more appropriate, *η* = *F*_*st*_/(1 − *F*_*st*_) can be used instead of *F*_*st*_ (Rousset, 1997). Under the infinite sites mutation model, pairwise *F*_*st*_ can be expressed in terms of the mean number of differences between sampled DNA sequences, called Hudson’s *F*_*st*_ estimator (Hudson et al., 1992). Thus, given two demes A and B, we can estimate pairwise *F*_*st*_ as:

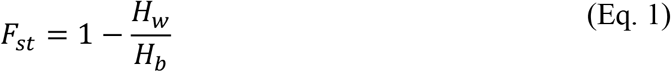

where *H*_*w*_ is the mean number of differences between pairs of DNA sequences sampled within the same deme, 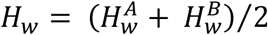, and *H*_b_ is the mean number of differences between pairs of DNA sequences sampled in different demes.

Slatkin, (1991) has shown that *F*_*st*_ can be expressed in terms of average coalescence times for a large variety of structured models, including the toroidal 2D stepping-stone, one type of spatially structured population model. According to the derivation of Slatkin, (1991), pairwise *F*_*st*_ for samples taken from demes A and B, at *i* and *j* steps apart along the *x* (e.g., longitude) and *y* (e.g., latitude) axes, respectively, can be expressed as:

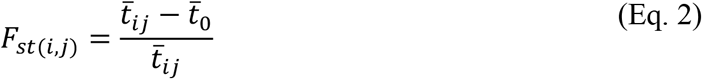

where 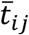 is the average coalescence time for two alleles sampled from a pair of demes at *i* and *j* steps apart, and 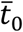 is the average coalescence time for two alleles sampled within the same deme. The equivalence between *Eq. 1* and *Eq. 2* becomes clearer if we consider that, under the assumption of small per-site mutation rate (*μ*), the expected number of differences between alleles, 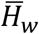 and 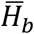, can be expressed in terms of average coalescence times (Slatkin, 1991). In particular, 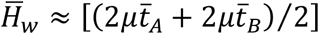 and 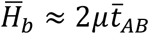, where 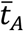 and 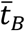 are the average coalescence times for two alleles randomly sampled within deme A and B, respectively, while 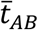 is the average coalescence times for two alleles sampled in A and B (Slatkin and Hudson, 1991). Thus, 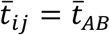 and 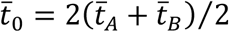. Assuming that deme size is homogeneous across the toroidal 2D stepping-stone model, then 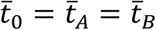.

In the case of a finite 2D stepping-stone model, it has been shown that, under a general model of dispersal, 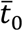 is independent from dispersal (*m*) and it can be estimated by only considering the total number of individuals in the population, 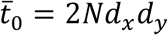, where *N* is the deme size and *d*_*x*_ or *d*_*y*_ are the number of demes along *x* and *y* axes, respectively (Strobeck, 1987; Hey, 1991). Instead, 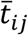 can be derived by dividing the coalescence process in two phases: *i)* the average amount of time required for two alleles sampled *i* and *j* steps apart to be in the same deme (*S*_*ij*_); and *ii)* the average time to coalesce once the two alleles occupy the same deme 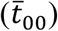 (Slatkin, 1991). Accordingly, 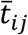 can be expressed as 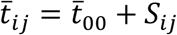. Then:

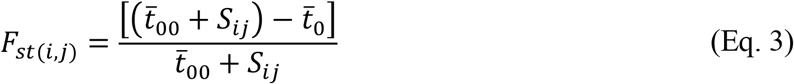

Given that the model assumes a homogenous deme size, 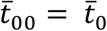 and therefore 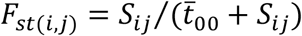. According to Slatkin, (2005), the term *S*_*ij*_ is defined as:

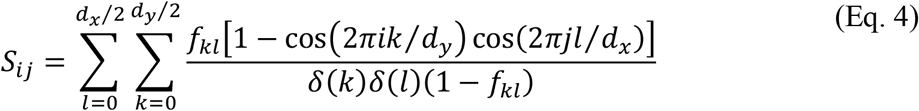

where *f*_*kl*_ = [1 − *m*(1 − cos(2π*k*/*d*_*y*_))]^2^ [1 − *m*(1 − cos(2π*l*/*d*_*x*_))]^2^ or *f*_*kl*_ = 0 if *k* = 0 and *l* = 0. The coefficients *δ*(*k*) = 1 or *δ*(*l*) = 1 if *k* = 0 or *l* = 0, respectively, and *δ*(*k*) = 1/2 or *δ*(*l*) = 1/2 if *k* > 0 or *l* > 0. As pointed out by Slatkin, (2005), this expression is a correction for typographical errors of *Eq. 8b* in Slatkin, (2003) and of *Eq. A9* in Slatkin, (1991).

In the context of HL&F, that is when a continuous habitat is suddenly broken apart in several habitat fragments, among which gene-flow is completely impaired, the expression for *F*_*st*(*i,j*)_ needs to be modified. To address this problem, we used an approach similar to the one proposed in Duforet-Frebourg and Slatkin, (2016) for modelling the expected coalescence time 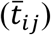 and *F*_*st*(*i,j*)_ for two alleles sampled at different times, let’s say in the present (modern DNA) and in the ancient past (ancient DNA). Our model assumes an initial 2D torus at dispersal-mutation-drift equilibrium, containing *d*_*x*_ x *d*_*y*_ demes, which undergoes HL&F at time *t*_*_, such that individuals occurring in each isolated habitat fragment can no longer disperse beyond the boundaries of the defined fragment. For simplicity, we assume that HL&F generates two habitat fragments, A and B, with 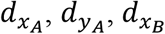 and 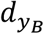 demes along the *x* and *y* axes. To derive an expression for *F*_*st*(*i,j*)_ in terms of expected coalescence times, we need to describe the properties of the coalescent process in the context of HL&F. It is useful then to think backward in time, considering that HL&F has occurred *t*_*_ generations in the past and that alleles are sampled in the present, *t* = 0 (see Supplementary Appendix S1: Figure S1 for a graphical visualization of the model). In particular, we decompose the coalescent process for two alleles sampled at *i* and *j* steps apart 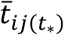 in three phases: *i*) the possible paths travelled since *t*_*_ by the ancestral lineage of each sampled allele across the available demes within the corresponding habitat fragment, 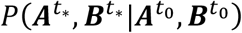; *ii*) the amount of time required for the two ancestral lineages to occupy the same deme in the continuous habitat, given that they were at *i* and *j* steps apart at the time of HL&F 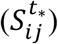; and *iii*) the average time to coalesce once the two ancestral lineages occupy the same deme in the continuous habitat 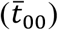. Thus, the expression for 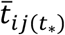 is:

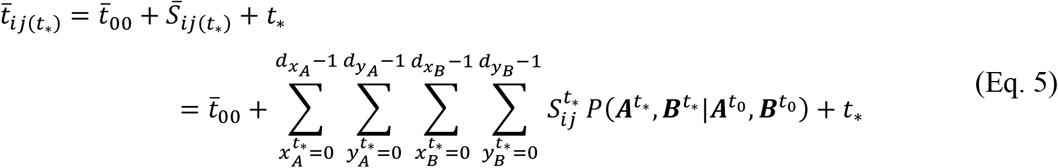

where, for habitat fragments A and B, 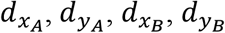 are the number of demes along the *x* and *y* axes, and 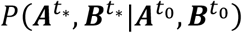 is the probability that two alleles sampled at 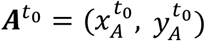 and at 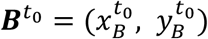 were at position 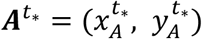 and 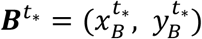 at the time of HL&F (*t*_*_). Such that:

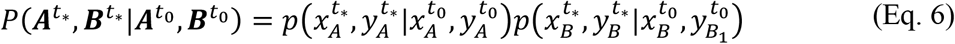

Using the method detailed in Duforet-Frebourg and Slatkin, (2016), we show how to compute 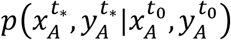, which procedure is valid also for 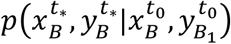. We consider a discrete random walk in a 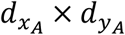 torus-like space with migration matrix *M* of size 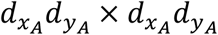:

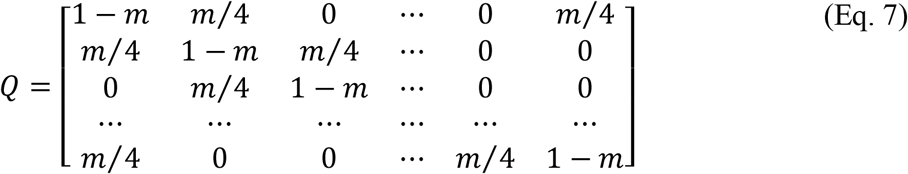

with rows and columns indices corresponding to the coordinates of each deme in the torus. The entries of the migration matrix indicate the probability of moving from one deme to another in one unit of time, which is 1 − *m* if the lineage was at the same location in the previous generation and *m*/4 if it was in any of the four adjacent demes. Then, using classical results on Markov chains, 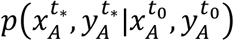 is obtained by taking the element 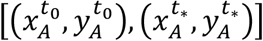 from the time *t*_*_ convolution of the migration matrix (the power *t*_*_ of the matrix *Q*). The term 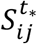 of *Eq. 5* is calculated from *Eq. 4* by defining 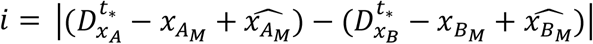 and 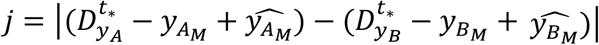. Assuming that deme (*x*_*A*_ = 0, *y*_*A*_ = 0) and (*x*_*B*_ = 0, *y*_*B*_ = 0) are the upper left corners of each ‘imaginary’ 2D plane fragment, 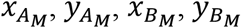 are the relative coordinates of the central demes, while the circumflexed symbol defines the respective coordinate projected in the continuous habitat, before HL&F. For instance, consider an initial 2D torus containing 78 × 78 demes, equivalent to a 40 × 40 2D plane. The 2D torus undergoes HL&F at time *t*_*_, resulting in two fragments A and B with 24 × 24 demes, which is equivalent to a 2D plane with 13 × 13 demes. Then the central deme within fragment A would have coordinates 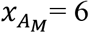 and 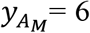, and can be projected to any coordinate in the initial continuous 2D torus, as long as the fragment size does not overpass the boundaries of the equivalent continuous 2D plane and does not overlap with any other fragment in the landscape, after HL&F. The 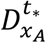 term indicates the shortest distance between 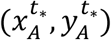 and (*x* = 0, *y* = 0) in the 2D torus-like space.

So far, the expressions used to derive 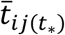 have referred to the consequences of *habitat fragmentation per se* (or disruption of gene-flow) on average coalescence times. To include *habitat loss*, corresponding to the reduction in available habitat within each fragment, we need to model 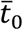 with an expression at non-equilibrium 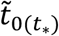. We used an approach similar to the one used in Maruyama, (1970) on the probability of identity by descent (Eq. 5-3) and to the expressions in Austerlitz et al., (1997) on the distribution of mean coalescence time in a population undergoing range expansion (Eq. 20). Consider a 2D torus composed by *d* × *d* demes, where, at each generation, individuals have a probability 1 − *m*_2*D*_ to remain in the same deme and a probability *m*_2*D*_/4 to disperse to each of the four adjacent demes (*m*_2*D*_/4 × 4 demes = *m*_2*D*_). Since a torus corresponds to the product of two 1D circular stepping-stones, one for each *x-* and *y*-axis, it is useful to describe the probability distribution of the migration patterns of two sampled lineages in a 1D circular stepping-stone. Such probability distribution is equivalent for both *x*- and *y*-axis, assuming that dispersal rate is equal in the two directions (isotropic dispersal). Thus, let ***M*** be a *d* × *d* circular matrix with elements *M*_*i,j*_, for *i, j* = 0, …., *d* − 1, describing the probability that two lineages *A*_1_ and *A*_2_ sampled at distance *i* in the present time were at distance *j* in the previous generation. Defining *m* = *m*_2*D*_/2, note that the probability *M*_0,0_ that lineages *A*_1_ and *A*_2_ sampled in the same deme (*i* = 0) were also in the same deme (*j* = 0) in the previous generation is:

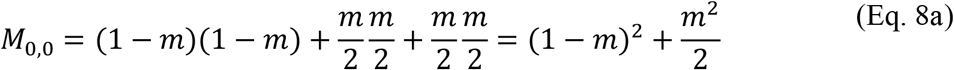

which is equivalent to the probability that none of the two lineages moved (1 − *m*)^2^, plus the probability that both moved clockwise (*m*/2)(*m*/2) plus the probability that both moved counterclockwise (*m*/2)(*m*/2). The probability *M*_0,1_ that lineages *A*_1_ and *A*_2_ sampled in the same deme, *i* = 0, were at distance *j* = 1 in the previous generation is:

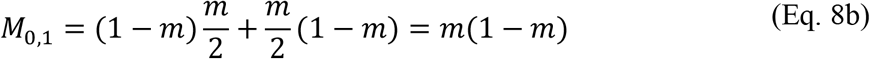

This is equivalent to the probability that one of the two lineages remained in the same deme in the previous generation (1 − *m*) times the probability that the other lineage moved clockwise (*m*/2), plus the probability that one of the two lineages remained in the same deme (1 − *m*) times the probability that the other lineage moved counterclockwise (*m*/2). Finally, consider *M*_0,2_ as the probability that lineages *A*_1_ and *A*_2_ sampled in the same deme, *i* = 0, were at distance *j* = 2 in the previous generation:

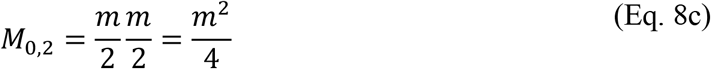

which is equivalent to the probability that in the previous generation one of the two lineages moved clockwise (*m*/2), while the other lineage moved counterclockwise (*m*/2). As an example, consider a 1D circular stepping-stone with *d* = 5, then the corresponding matrix ***M*** would be:

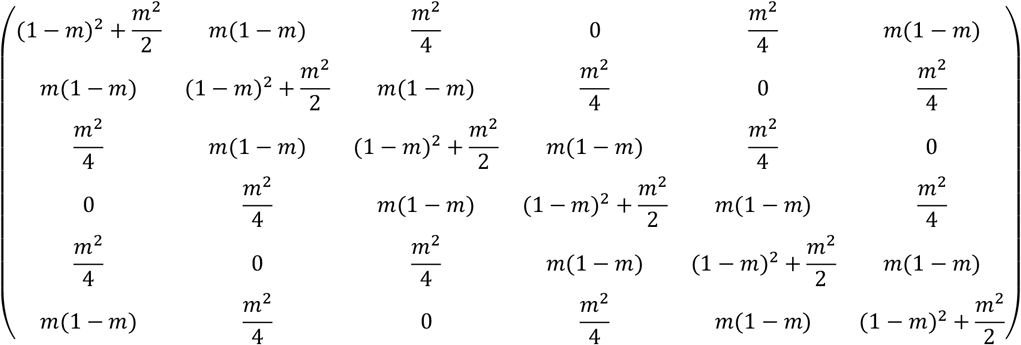

Given the matrix ***M***, it is possible to compute the mean coalescence time at the next generation ***T***^′^ as:

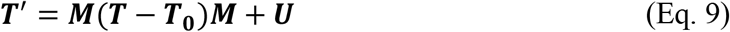

where ***T*** is a *d*x*d* matrix of mean coalescence times of lineages initially separated by a distance *i* and *j* on the *x*- and *y*-direction, respectively; ***T***_***0***_ is a *d*x*d* matrix with 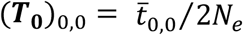 and 0 elsewhere; and ***U*** is a *d*x*d* matrix whose elements are 1 everywhere. The expression ***T*** − ***T***_***0***_ takes into account that if two lineages were in the same deme in the previous generation, then the contribution of 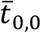 needs to be weighted by the probability of not coalescing (1 − 1/2*N*). In fact, at *i* = 0 and *j* = 0, (***T***)_0,0_ − (***T***_***0***_)_0,0_ takes the form 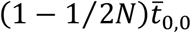. Moreover, note that in *Eq. 9* the matrix ***M*** is multiplied at the left and at the right of ***T*** − ***T***_***0***_, since it is necessary to consider the possible migration patterns in both *x*- and *y*-axis. Ultimately, the matrix ***U*** is used to add one time unit at each generation, given that lineages did not coalesce. At any generation after HL&F, the mean coalescence time for pairs of lineages at distance *i* and *j* can then be calculated by iterating *Eq. 9*.

There are now all the necessary elements to re-rewrite *Eq. 1, 2* for a model of population divergence in space and time, such that:

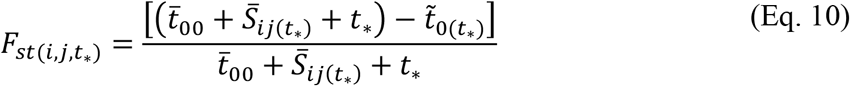

For a better interpretation of our results, we define 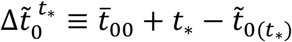 as the *drift-isolation term*, which determine the rate of ‘independent’ evolution of each habitat fragment due to *habitat loss* since HL&F at time *t*_*_. In particular, 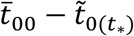 determines the rate of loss in genetic diversity within demes, while *t*_*_ considers the mutations that accumulate in each lineage in the two habitat fragments. To put it in another way, *t*_*_ corresponds also to the time is necessary to wait such that two alleles have the possibility to coalesce. This is because two lineages cannot coalesce until they occupy the same deme, condition that requires a continuous habitat where gene-flow is possible between the two sampled alleles at *i* and *j* steps apart. Thus, we can re-arrange *Fst*(*i,j,t*_*_) as:

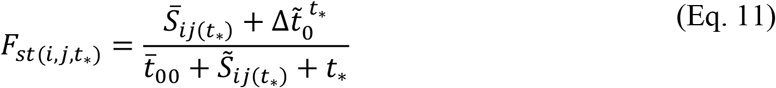

This last equation will turn out to be useful when we discuss the effect of HL&F on IBD pattern over time in the *Discussion* section.

## Results

### Instantaneous HL&F

In the present study, we investigated the effect of landscape-scale habitat loss and fragmentation on ‘isolation-by-distance’ patterns (IBD). Overall, our results suggest that significant IBD signatures can be detected for surprisingly long periods (on the order of thousands of generations) after HL&F. We showed that dispersal rate (0.02 – 0.1) and sampling strategy (*local* vs *random*) influence the time of loss of IBD (*TIBD*) under the *instantaneous HL&F* scenario (Fig. 1b). On average, ‘isolation-by-distance’ pattern persists for longer time at lower dispersal rates (*m* = 0.02, *T*_*IBD*−*local*_ ≈ 700; *m* = 0.1, *T*_*IBD*−*local*_ ≈ 50; Fig. 1b, Supplementary Appendix S1: Figure S2a). The influence of dispersal rate on *T*_*IBD*_ becomes more noticeable when alleles are randomly sampled within a habitat fragment (*random*) compared to a within-deme (*local*) sampling scheme (*m* = 0.02, *T*_*IBD*−*random*_ ≈ 1500; *m* = 0.1, *T*_*IBD*−*random*_ ≈ 200). Furthermore, this suggests that sampling strategy (*local* vs *random*) can significantly influence *T*_*IBD*_. Larger absolute differences between *T*_*IBD*−*local*_ and *T*_*IBD*−*random*_ were identifiable at lower dispersal rate. For instance, at *m* = 0.02, we estimated an average *T*_*IBD*−*local*_ of 700 and a *T*_*IBD*−*random*_ of 1500 generations (*m* = 0.02; U = 1974, p-val<0.001), whereas at *m* = 0.1, *T*_*IBD*−*local*_ was detected at 50 and *T*_*IBD*−*random*_ at 200 generations (*m* = 0.1; U = 2969.5, p-val<0.001). These results were consistent when IBD patterns were estimated using Wright’s F_st_ instead of Slatkin’s R_st_ (Supplementary Appendix S1: Figure S3). Carrying capacity and mutation rate appear to influence *T*_*IBD*_ much less than dispersal rate, for both *local* and *random* sampling schemes (Fig. 1c, d). The distribution of *T*_*IBD*_ for each independent simulation (Fig. 1c, Supplementary Appendix S1: Figure S2b, e) and average IBD p-value (Fig. 1c) are nearly indistinguishable among the three values of carrying capacity herein tested (50 – 200). Small differences were observed in R_st_-based IBD pattern between *K* = 200 and *K* = 50/100, but they were nearly absent when IBD was estimated from F_st_ (Supplementary Appendix S1: Figure S4). With some exceptions, we did not observe differences in *T*_*IBD*_ in relation to mutation rate (Fig. 1d). In fact, for mutation rates between 10^−2^ and 10^−5^ per locus per generation, our results show no difference in the time at which IBD tends to be lost. Different is the case for mutation rate equal to 5 × 10^−6^, since IBD tends to be lost earlier than the other tested values. The independence of *T*_*IBD*_ from mutation rate is further supported by the lack of differences in the temporal dynamic of IBD slope among simulations with different values of mutation rate (Supplementary Appendix S1: Figure S2c, f).

The number of genotyped markers has a major role in determining the time of IBD loss (Supplementary Appendix S1: Figure S6). As an example, when we simulated an *instantaneous HL&F* scenario (*K* = 50; *m* = 0.04; *μ* = 5 × 10^−4^) and sampled either 15, 30, 100 or 400 microsatellite loci, we found that the estimated time before the loss of a significant IBD signature (*T*_*IBD*_) correlated with the number of genotyped markers. We did not observe major differences in *T*_*IBD*_ for 15 and 30 loci (U_15-30_ = 193.5, p-value = 0.003), but strong discordance between *T*_*IBD*_ for 15/30 loci and 100/400 loci (U_15-100_ = 597.5, p-value<0.001; U_30-400_ = 105.5, p-value<0.001). Similar results were obtained for IBD patterns measured using Wright’s F_*s*t_; however, in this case, differences of *T*_*IBD*_ were less strong (U_15-100_ = 797, p-value<0.001), but still important (Supplementary Appendix S1: Figure S7). A qualitative comparison with the previous results suggests that the number of genotyped loci can have a greater impact on *T*_*IBD*_ than dispersal rate (e.g., Supplementary Appendix S1: Figure S3, S7).

### Range expansion prior to instantaneous HL&F

We also considered a more complex scenario where *instantaneous HL&F* occurs after a range expansion, i.e., in a non-equilibrium population (Fig. 2a). Thus, two evolutionary processes are expected to contribute to the observed spatial genetic pattern: *range expansion* and *HL&F*. Several *range expansion + instantaneous HL&F* scenarios were simulated, where HL&F occurred at different time points since the onset of the range expansion (*T*_*exp*_ = 600, 750, 900, 1050 and 1500). We observed a significant increase in *T*_*IBD*_ for the *range expansion + instantaneous HL&F* compared to the *instantaneous HL&F* scenario (Fig. 2a, Supplementary Appendix S1: Figure S8a, c). Such differences were observable with both *local* and *random* sampling, but more noticeable with the *random* sampling (*equilibrium* vs *T*_*exp*_ = 600: U_*local*_ = 1740.5, p-value = 0.01; U_*random*_ = 1136, p-value < 0.001). Differences between *T*_*IBD*−*equilibrium*_ and *T*_*IBD*−*expansion*_ were larger for recent than for ancient range expansions (*equilibrium* vs *T*_*exp*_ = 900: U_*random*_ = 1200.5, p-value = 0.027), converging to the behavior of the *instantaneous HL&F* for more ancient range expansions (*equilibrium* vs *T*_*exp*_ = 1500: U_*random*_ = 1244.5, p-value = 0.8774). Similar results were obtained for IBD-patterns estimated using Wright ‘s F_st_ (Supplementary Appendix S1: Figure S9). The temporal dynamics of pairwise R_st_ indicates that differences between *T*_*IBD*−*equilibrium*_ and *T*_*IBD*−*expansion*_ are related to the larger effect of geographical distance on pairwise R_st_ in the *range expansion + instantaneous HL&F* compared to the *instantaneous HL&F* scenario. For instance, using relative weights analysis, we estimated a relative rescaled weight for geography on pairwise R_st_ (*RRW*_*geography*_) of ≈ 1% for *equilibrium* and ≈ 7% for *T*_*exp*_= 600 scenarios (Supplementary Appendix S1: Figure S11a, b). The results show that *range expansion* has also the effect of increasing the variance around the mean R_st_ (shaded area in Supplementary Appendix S1: Figure S11a). As shown by the behavior of *T*_*IBD*−*expansion*_, discrepancies in pairwise R_st_ (or F_st_) between the *range expansion + instantaneous HL&F* and the *instantaneous HL&F* scenario decrease with the age of the range expansion (*RRW*_*geography*_ ≈ 1% for *equilibrium*; *RRW*_*geography*_ ≈ 1.5% for *T*_*exp*_= 1500; Supplementary Appendix S1: Figure S11a).

### Two-steps HL&F

We assessed the behavior of the time of loss of a significant IBD signature (*T*_*IBD*_) in a scenario where two consecutive HL&F events generated the same HL&F landscape configuration as in the *instantaneous HL&F* (Fig. 2b). We tested the effect of the time since the first HL&F event (*T*_*split*_) on *T*_*IBD*_ and compared it to the *T*_*IBD*_ in an equivalent *instantaneous HL&F* scenario (*T*_*equilibrium*_). We observed that *T*_*IBD*_ increased with *T*_*split*_. For instance, we observed a difference in *T*_*IBD*_ of ∼ 400 generations between *T*_*equilibrium*_ and *T*_*split*_ = 500 scenarios and a difference of ∼ 2000 generations between *T*_*equilibrium*_ and *T*_*split*_ = 1500. Similar results are reported also for Wright ‘s F_st_ – IBD pattern (Supplementary Appendix S1: Figure S10). As for the other investigated scenarios, the effect of *Two-steps HL&F* is even more marked for *random* than *local* sampling. Pairwise R_st_ averaged across simulations or pairwise R_st_ for single simulation replicates (Supplementary Appendix S1: Figure S11c, d) shows that the longer persistence of IBD in the *Two-steps HL&F* scenario, compared to the *instantaneous HL&F* scenario (*equilibrium*), is related to the larger effect of geographic distance on R_st_ (*RRW*_*geography*_ ≈ 1% for *equilibrium*; *RRW*_*geography*_ ≈ 8% for *T*_*split*_= 1500).

### Population divergence in space and time

F_st_ between pairs of populations can be expressed in terms of expected coalescence time and can be calculated for various types of structured model, such as a 2D-stepping stone model (Slatkin, 1991). Using the approach developed in Duforet-Frebourg and Slatkin, (2016), we proposed an analytical model of isolation-by-distance in time in the context of habitat loss and fragmentation. The proposed model was used to aske the following questions: ‘how does genetic differentiation accumulate over time between two habitat fragments after HL&F?’ Does geography contribute to the level of genetic differentiation measured between habitat fragments for long time? As outlined in *Materials and Methods* (*Eq. 11*), genetic differentiation over time between two isolated habitat fragments can be expressed as: 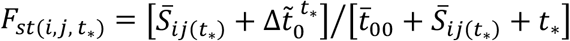, where 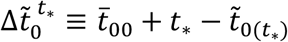. For the sake of clarity, we will refer to 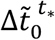 as the *drift-isolation term* and to 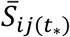 as the *spatial term*. In Fig. 3-4, we represent the behavior of 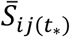 and 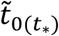 in relation to dispersal rate. In particular, Fig. 3a shows the impact of dispersal rate on the relative distribution of *S*_*ij*_ values within habitat fragment 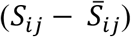, highlighting larger differences in *S*_*ij*_ among demes when dispersal rate is low (scaled standard-deviation *σ*_*m*=0.02_ = 8,715 *vs σ*_*m*=0.1_ = 1,742). In Fig. 3b-c, we show the probability 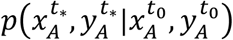 that a lineage sampled at time *t* = 0 in the center of a 9 × 9 plane habitat fragment (equivalent to a 16 × 16 torus) was originally at any other position within the fragment, respectively, at *t*_*_ = 50 and *t*_*_ = 1000. As expected, higher is dispersal rate, higher is the probability that the lineage sampled in the center of the fragment was further apart at the time of HL&F (Fig. 3b). The same is true for the time of HL&F event. Older is the HL&F event, higher is the probability that the lineage sampled in the center of the fragment was further apart at the time of HL&F (*t*_*_ = 50, Fig. 3b *versus t*_*_ = 1000, Fig. 3c).

**Figure 3.**
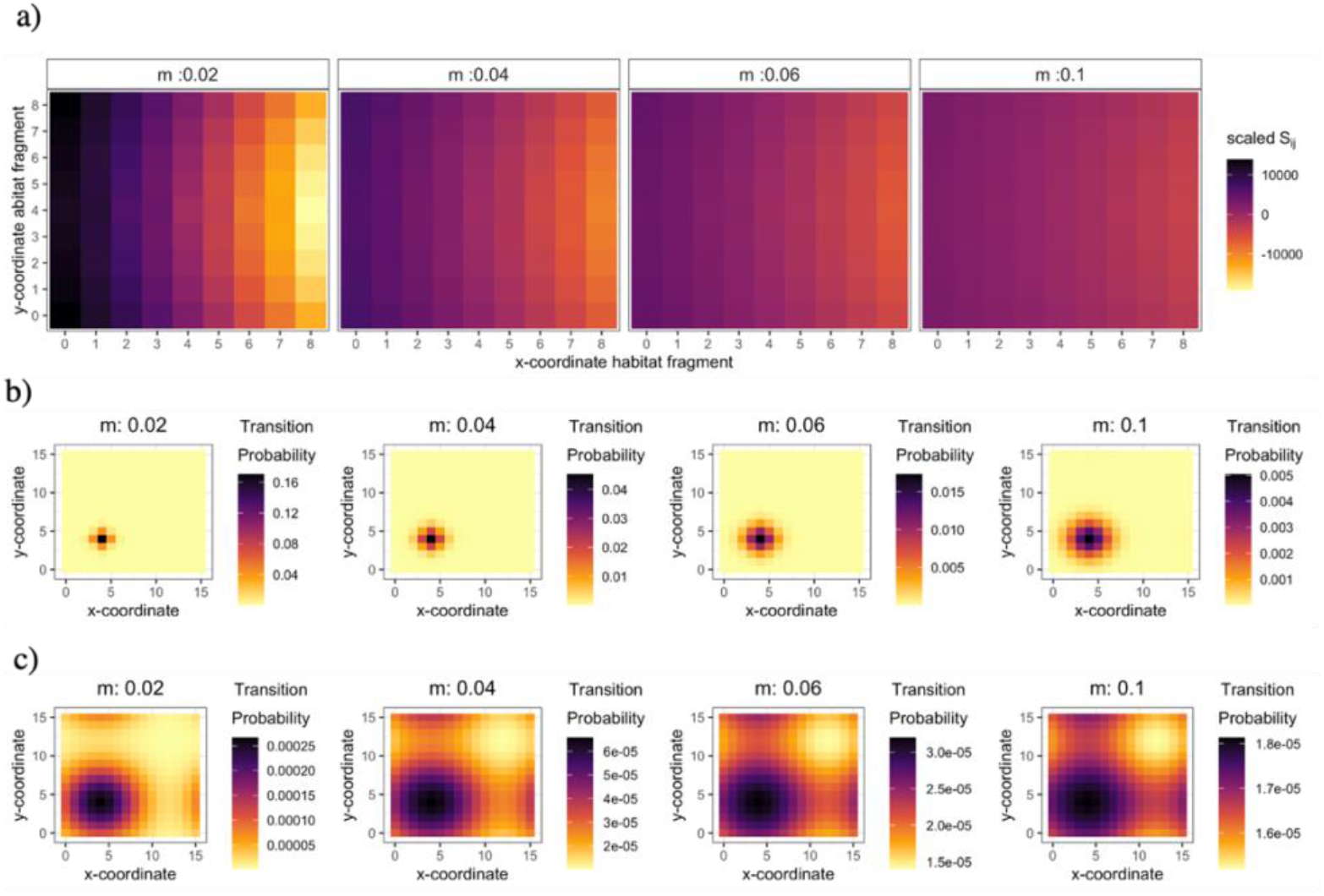
Behaviour of the spatial term 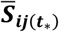 as function of geographical position and time since HL&F (t_*_). **a)** Relative distribution of *S*_*ij*_ for all *x*_*A*_ and *y*_*A*_ within fragment A (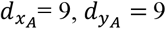 in a plane and 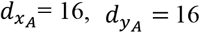 in a torus) by fixing the sampled location of a random allele in fragment B (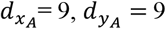 in a plane) at *x*_*B*_= 4 and *y*_*B*_= 4 (centre of 9 × 9 plane). Absolute *S*_*ij*_ values for each dispersal rate were scaled with respect to its mean *S*_*ij*_ across the fragment, e.g., *S*_*ij*(*m*:0.02)_ − *mean*(*S*_*ij*(*m*:0.02)_). Transition probabilities 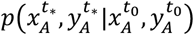 that a lineage sampled at time *t* = 0 in the center of fragment A was at any other position within the fragment A, respectively, at **b)** *t*_*_ = 50 and **c)** *t*_*_ = 1000, as function of dispersal rate.

**Figure 4.**
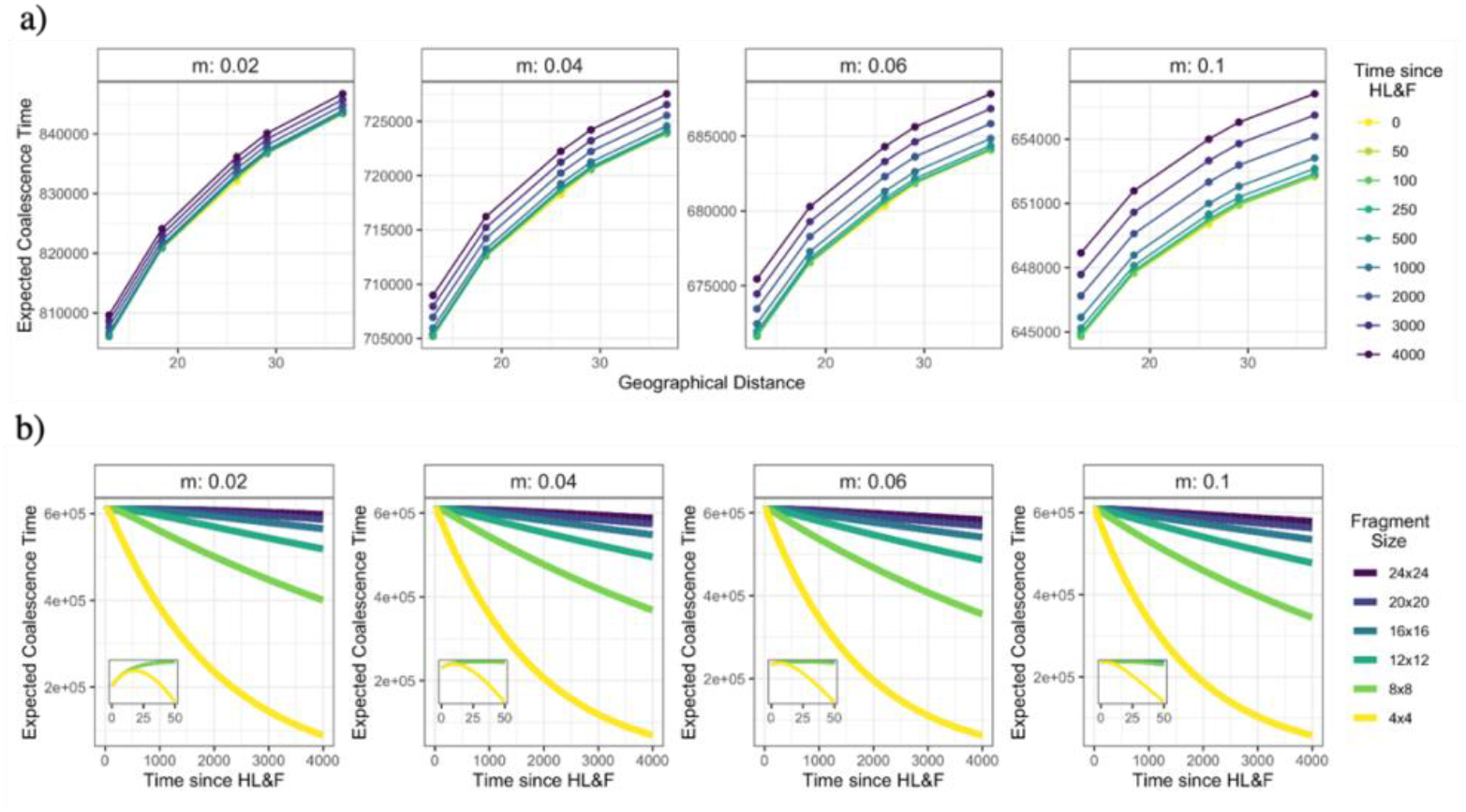
Non-equilibrium mean coalescence time between and within habitat fragments. **a)** Change in mean coalescence time between two alleles sampled at the centre of two isolated habitat fragments, 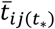, as function of dispersal rate and geographical distance (*Eq. 5*). **b)** Decay in mean coalescence time for two alleles randomly sampled within the same deme in a 78 × 78 torus-like space undergoing habitat contraction 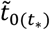. The results of a) and b) show i) a stronger effect of time since HL&F on 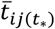 when dispersal rate is high, and ii) a slower decay in 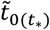 at low than high dispersal rate. Numerical results were obtained assuming a 78×78 torus (equivalent to a 40×40 plane) before HL&F and five 16 × 16 torus-like fragments (equivalent to a 9×9 plane) after HL&F.

Fig. 4a show the dependence of 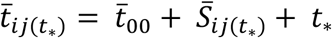 on geographical distance, time since HL&F (*t*_*_) and dispersal rate. We found that lower is the dispersal rate, lower is the relative impact of time since HL&F (*RRW*_*t*\*_) compared to geography (*RRW*_*geography*_) on pairwise *F*_*st*(*i,j, t*\*_) (*m* = 0.02, *RRW*_*geography*_ ≈ 99%, *RRW*_*t*_* ≈ 1%; *m* = 0.1, *RRW*_*geography*_ ≈ 80%, *RRW*_*t*_* ≈ 20%). In Fig. 4b we show the decay in 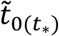 after HL&F as a function of habitat fragment size and dispersal rate. As expected, smaller is the habitat fragment size after HL&F, steeper is the decay in expected coalescence time. Moreover, lower is dispersal rate, slower is the decay in expected coalescence time.

Using the expressions describing the decay in 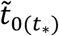 and the increase in 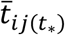, we can numerically compute 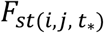. The results are displayed in Fig. 5-6, which shows how 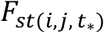 changes over time, as function of geographical distance, dispersal rate, deme population size and habitat loss. For the sake of clarity, we decided to represent only two extreme cases among those here considered, namely a HL&F event that break a 78 × 78 torus-like space in fragments of either 24 × 24 or 4 × 4 demes. Regarding dispersal rate, relative weights analyses showed that the relative contribution of geography on 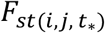 decreases with increasing dispersal rate. For instance, in the case of habitat fragment size 24 × 24, we quantified a 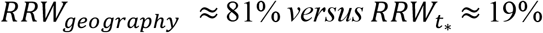 for the lower dispersal rate 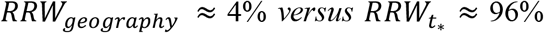 for the higher dispersal rate (*m* = 0.1). Concerning deme population size, the relative contribution of geography on 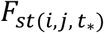 remains nearly constant with increasing deme population size. For habitat fragment size 24 × 24, we quantified a 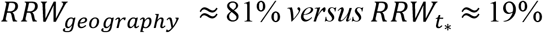 for the lower deme population size (*N* = 50) and a 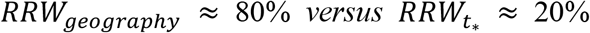 for the higher deme population size (*N* = 200). In Fig. 5-6, we also show that the relative contribution of geography on 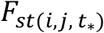 strongly depends on habitat loss. Accordingly, relative weights analysis quantified, for a smaller habitat fragment size (4 × 4), a 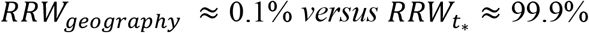 at low dispersal rate (*m* = 0.02) and a 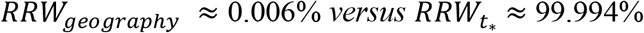 at high dispersal rate (*m* = 0.1). A wider exploration of the parameter space is shown in Fig. 7, where *RRW*_*geography*_ is displayed for any combination of deme population size (*N* = 25 − 300), dispersal rate (*m* = 0.002 − 0.1) and habitat loss (24×24, 20×20, 16×16, 12×12, 8×8, 4×4). The results show that dispersal rate and habitat loss are the factors that consistently determine the contribution of geography on pairwise F_st_.

**Figure 5.**
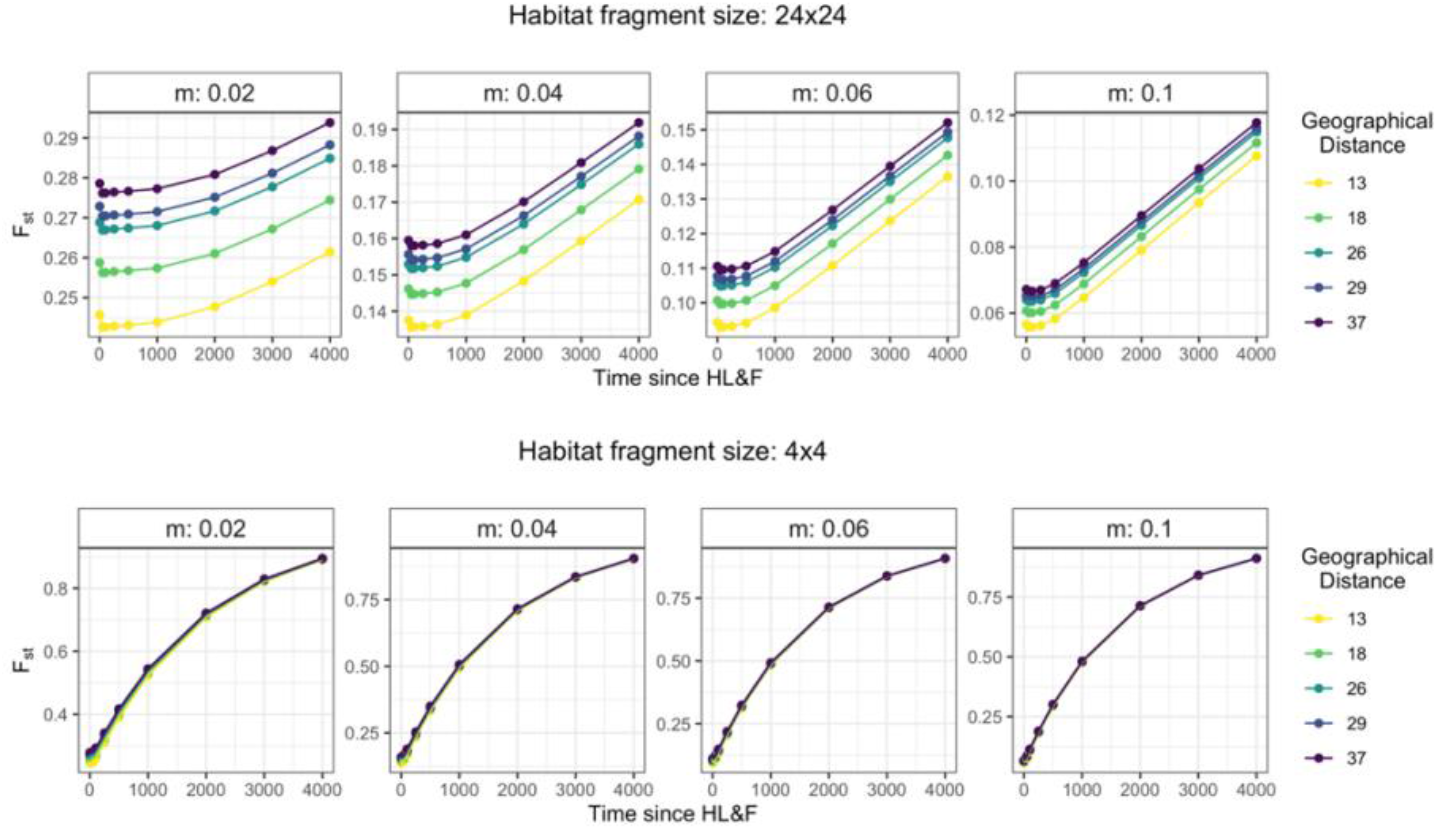
The effect of dispersal rate on expected pairwise *F*_***st***_ in a torus-like space undergoing HL&F. First and second rows refer to results for habitat fragments of size 24×24 and 4×4 (torus-like fragments), respectively. Expected pairwise *F*_*st*_ were computed from *Eq. 10*. The analytical model suggests that time since HL&F has a stronger influence on pairwise *F*_*st*_ at high dispersal rate and/or when habitat undergoes severe habitat loss (4 × 4).

**Figure 6.**
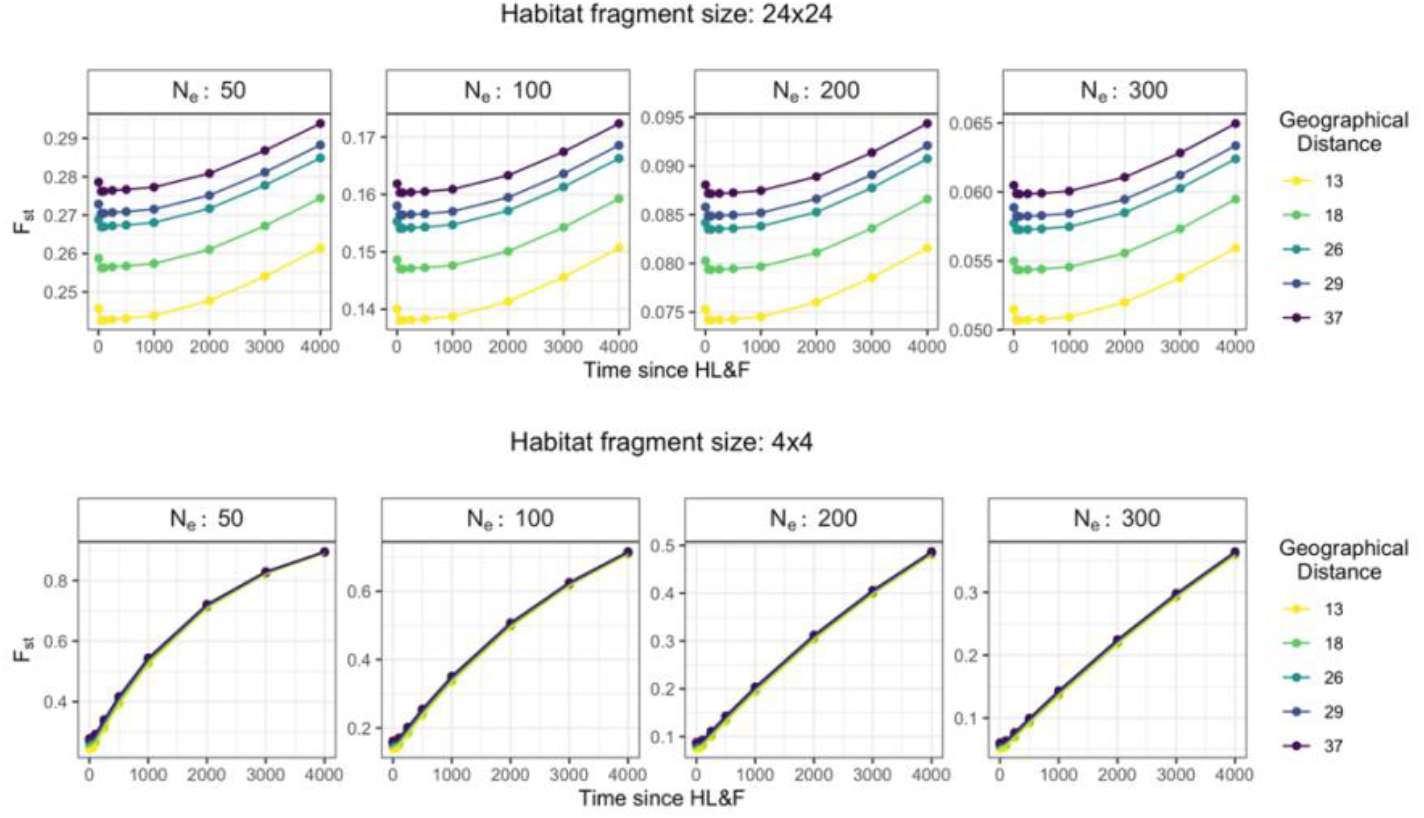
The effect of deme population size on expected pairwise *F*_*st*_ in a torus-like space undergoing HL&F. First and second rows refer to results for habitat fragments of size 24 × 24 and 4 × 4 (torus-like fragments), respectively. Expected pairwise *F*_*st*_ were computed from *Eq. 10*. The analytical model suggests that the relative contribution of geography on pairwise *F*_*st*_ is little affected by deme population size.

**Figure 7.**
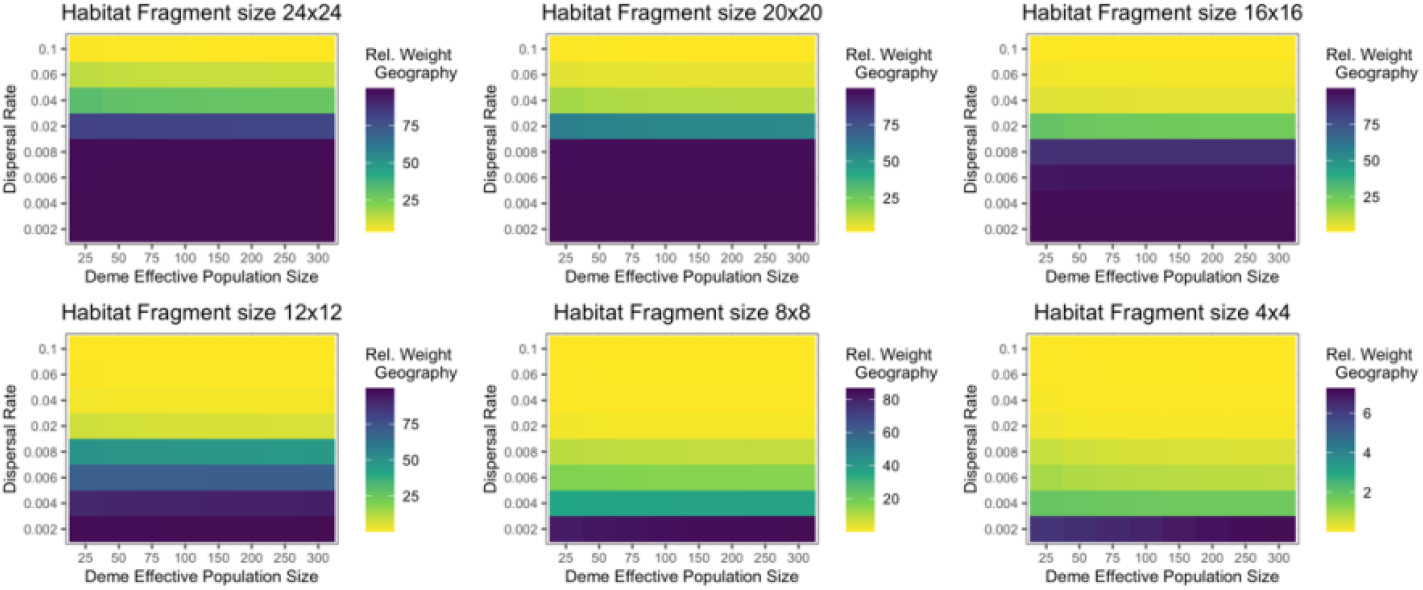
The effect of dispersal rate, deme population size and habitat loss on the relative contribution of geography on expected pairwise F_**st**_ in a torus-like space undergoing HL&F. Each panel refers to results obtained in a torus-like space of size 78×78 undergoing HL&F, leading to the formation of nine torus-like fragments of size 24×24, 20×20, 16×16, 12×12, 8×8 or 4×4. Expected pairwise F_st_ was numerically computed from *Eq. 10* for different time points after HL&F: 50, 100, 250, 500, 1000, 2000, 3000, 4000 generations. Rel. Weight Geography: relative contribution of geography computed using the relative weight analysis (see *Materials and Methods*). Overall, the results suggest that dispersal rate and habitat loss are the factors that determine the most the relative contribution of geography on pairwise F_st_.

## Discussion

### Overview of the study

Classical IBD models relies on the assumption that populations are at mutation– dispersal–drift equilibrium and that demes are connected by ongoing gene-flow. In the present study, we used individual-based spatially-explicit simulations to explore non-equilibrium isolation-by-distance patterns (IBD) during habitat loss and fragmentation (HL&F). In particular, we investigated two scenarios in which populations were at mutation–dispersal– drift equilibrium prior to HL&F (*instantaneous HL&F* and *Two-steps HL&F*) and one scenario were HL&F occur in a non-equilibrium population (*range expansion prior to HL&F*). We start considering the case of an *instantaneous HL&F*, assessing the influence of deme population size, dispersal and mutation rates on the time of IBD loss (*T*_*IBD*_). Since, natural populations may not necessarily be at mutation–dispersal–drift equilibrium, we also investigated a scenario in which an *instantaneous HL&F* occur after a *range expansion*. The last explored scenario focuses on the effect of *Two-steps HL&F* on IBD patterns. In fact, it is likely more realistic to consider that landscape-level HL&F occurs gradually over time following fine-scale spatial-temporal processes (Mandelbrot, 1982; Krummel et al., 1987; Milne, 1988; Solé and Manrubia, 1995).

The aim of the present study is to provide some insights on why significant IBD patterns can still be observed in fragmented natural populations (e.g., Schneider et al., 2010; Aleixo-Pais et al., 2019). The demographic history of species is certainly complex and the simple scenarios explored in this work do not represent a comprehensive understanding of non-equilibrium IBD pattern in the context of HL&F. However, we attempted to consider scenarios that are likely common in real populations and thus, may contribute to a better interpretation of non-equilibrium IBD pattern from wild populations. This is the case, for instance, for *range expansion + instantaneous HL&F* scenario. In fact, observed present-day genetic data suggest that range expansion has shaped the genetic diversity of many taxa across the world. A well-known example is given by species from Eurasia that have undergone expansion following the retreat of ice sheets at the end of the Last Glacial Period (LGM), about 20 thousand years ago (Hewitt, 2000, 1999; Lattin, 1967; Taberlet et al., 1998; Juřičková et al., 2014). We conclude the present work proposing a theoretical model of *isolation-by-distance in time* in the context of HL&F, which could be interpreted also as a model of *population divergence in space and time*. In *Materials and Methods*, we presented the details of the model by first summarizing the original theoretical results on a toroidal 2D stepping-stone model and then extending the theory in the context of habitat loss and fragmentation. Our analytical model shows that non-equilibrium *F*_*st*_ can be expressed as 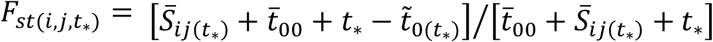. We have shown that the dependence of 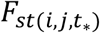 on geographical distance relies on the relative contribution of the *spatial term* 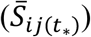 compared to the *drift-isolation term* 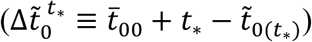. In the *spatial term*, the effect of geographical distance relies on how dispersal rate influences the *i*) expected coalescence time between two genes at *i* and *j* steps apart, in the continuous torus-like space, before HL&F; and *ii*) the probability that genes sampled at time *t* = 0, respectively, in fragment A and B were in other demes within each fragment at the time of HL&F (*Eq. 5*). In the *drift-isolation term*, geographical distance (related to habitat fragment size) and dispersal rate influence the decay in 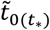, which occurs at a slower rate i) for low compared to high dispersal rate (Fig. 4b) and ii) for large compared to small fragment size (Fig. 5-6). To put it briefly, the numerical results have shown that dispersal rate and habitat fragment size are the two most important factors influencing the relative contribution of geography on pairwise *F*_*st*_ (Fig. 7). This short overview of the analytical model is going to be useful for discussing the simulation results presented below.

### Instantaneous HL&F

Overall, our simulation results have shown that IBD generated before HL&F can be maintained for thousands of generations after HL&F. We started investigating the contribution of deme population size, dispersal or mutation rate on the persistence of IBD during HL&F. We found that the time of IBD loss (*T*_*IBD*_) is mainly dependent on dispersal rate, and little affected by population size and mutation rate. In particular, we showed that at low dispersal rate IBD is maintained for a larger number of generations after HL&F compared to an equivalent scenario at higher dispersal rate (Fig. 1a). Conversely, when testing different values of either population size or mutation rate, *T*_*IBD*_ remained nearly unchanged across the different parameters combinations (Fig. 1b, c). We interpret these results using coalescent theory and the relationship between average coalescence times and pairwise *F*_*st*_ (*Eq. 11*). In our simulation results (Supplementary Appendix S1: Figure S6-S7), it is shown that, following HL&F, IBD patterns are lost due to the fact that pairwise *F*_*st*(*i,j*)_ are no longer distinguishable in relation to geographical distance. The analytical model suggests that this condition is satisfied when the relative contribution of 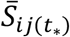 is much smaller than 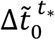, which can be due to either *i*) a very ancient HL&F (*t*_*_), or *ii*) to a large reduction in total population size within habitat fragment 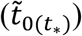; *iii*) or to both phenomena. The decrease in *T*_*IBD*_ with increasing dispersal rate observed in the simulation results can be understood from the relationship between *S*_*ij*_ and dispersal rate (Fig. 3a). In fact, higher is dispersal rate, lower is *S*_*ij*_, and thus a smaller amount of time (since HL&F) is required for the *spatial term* to become negligible compared to 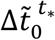 (Fig. 4a and 7).

The analytical model also explains why *T*_*IBD*_ is mainly influenced by dispersal rate, and little by mutation rate and population size. In fact, as shown by Slatkin, (1991), one of the advantages of expressing *F*_*st*_ in terms of coalescence times is that it does not depend on mutation rate (*μ*). This is analytically derived under the assumption that *μ* ≪ 1. Accordingly, our simulations showed that mutation rate does not influence *T*_*IBD*_. An exception was found for *μ* = 5 *x* 10^−6^, which can be explained by the fact that the simulated population did not have enough time before HL&F (10^4^ generations) to reach mutation-dispersal-drift equilibrium. In fact, it has been shown that in a 2D stepping-stone model, pairwise *F*_*st*_ approaches equilibrium over a timescale on the order of *μ*^−1^, which in this case would be 10^6^ generations (Slatkin and Barton, 1989; Barton et al., 2013). Both simulation and numerical results have shown that deme population size has little effect on *T*_*IBD*_ (Fig. 1b) and on the relative contribution of geography on pairwise *F*_*st*_ (Fig. 7). This can be understood by the fact that i) the *spatial* term 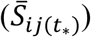 is independent from deme population size (*Eq. 4*), ii) the *drift-isolation* term 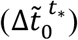 is smaller than the *spatial* term for any deme population size values, except under strong habitat loss and long time since HL&F (Supplementary Appendix S1: Figure S12), and that iii) the absolute decrease in 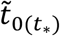 is nearly identical among different values of deme population size values, except under strong habitat loss (Supplementary Appendix S1: Figure S12).

Our simulation results have also shown that sampling scheme can contribute significantly to IBD, and thus to influence *T*_*IBD*_, when dispersal rate is relatively low (Fig. 1 and 2). In particular, we show that a scattered sampling within habitat fragments (*random*) contribute in maintaining IBD for longer after HL&F than a localized sampling (*local*). The effect of sampling scheme on IBD was consistent among all HL&F scenarios here simulated, thus we give an interpretation of this result in this section, which will be valid for the other HL&F scenarios. In particular, we suggest that the longer *T*_*IBD*_ in *random* sampling is due to the higher expected coalescence times for *random* than *local* sampling scheme, as shown by measuring expected heterozygosity over time (Supplementary Appendix S1: Fig. S13). Referring to *Eq. 10*, this means that 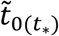 will be higher in *random* than *local* sampling, resulting in a lower contribution of the *drift-isolation term* on pairwise *F*_*st*_. Instead, *S*_*ij*_ is expected to remain nearly unchanged between the two sampling schemes, given that *random* sampling collects genetic data with equal probability from the available demes within habitat fragment. Therefore, changes in the temporal dynamic of *F*_*st*(*i,j*)_ for the two-sampling scheme are mostly related to differences in the expected coalescence time for the alleles sampled within fragments 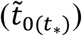.

### Range Expansion before Instantaneous HL&F

Range expansion is a common process in nature (Hewitt, 1999; Lawson Handley et al., 2011), which has shaped spatial genetic patterns in several taxa across the world. It is characterized by successive events of colonization that reduce genetic diversity towards the direction of expansion (e.g., Austerlitz et al., 1997; Edmonds et al., 2004; Hallatschek and Nelson, 2008). A mechanism that can explain such decay in genetic diversity is called ‘gene-surfing’, which refers to the random spread of rare alleles that, by chance, can reach very high frequencies at the front of the expansion (Klopfstein et al., 2006). The non-homogenous spatial distribution of allele frequency due to a recent range expansion can generate large differences in allele frequency between the origin and the front-wave of the spatial expansion. This can in turn lead to higher genetic differentiation (e.g., *F*_*st*_) compared to an equivalent model at mutation-dispersal-drift equilibrium (Hallatschek et al., 2007). The consequences of range expansion on the spatial distribution of allele frequencies point out that both geographically-limited dispersal and range expansion can generate isolation-by-distance patterns. For instance, Meirmans, (2012) found through spatially-explicit simulations that in a colonization scenario from a single refugium, the percentage of significant tests in IBD were higher (44%) than in an equilibrium scenario (24%). These series of studies led us to consider that range expansion may probably contribute to the persistence of IBD pattern after HL&F. Earlier simulation results by Mona et al., (2014) have, in fact, shown that under a scenario of *range expansion in a fragmented habitat, F*_*st*_ average and variance are higher than i) a non-fragmented habitat, and an equivalent fragmented population that has not undergone range expansion. As expected, the strength of such discrepancy between equilibrium (*eq*) and expanded (*exp*) populations decreases with the age of the range expansion 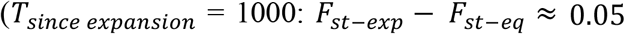 and 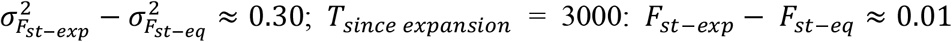 and 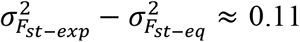 in Mona et al., 2014). In the present study we examined a different scenario, in which *habitat loss and fragmentation occurs after range expansion* (Fig. 2a). Similar to Mona et al., (2014), we have studied the impact of the age of the range expansion on the persistence of IBD patterns after HL&F. Our results suggest a significant effect of recent range expansion on IBD patterns, for instance, by increasing *T*_*IBD*_ of about 1000 generations in *T*_*exp*_ = 600 compared to an equilibrium model (Fig. 2a, Supplementary Appendix S1: Figure S8a, c, Figure S9). However, the influence of range expansion on *T*_*IBD*_ decreases with the age of the onset of the expansion, converging to the values of an equivalent *instantaneous HL&F*. As for the instantaneous HL&F scenario, we interpret the simulation results in terms of expected coalescence time. Following the theoretical results derived in Austerlitz et al., (1997) on the impact of colonization on coalescence times, within-deme coalescence times are expected to be lower in a colonizing population than in an equivalent population at equilibrium. Consequently, the average coalescence time for two alleles sampled within the same deme 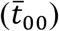 is expected to be lower than an equilibrium population. Accordingly, also 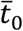 will be lower than expected. However, in this case 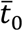 is not the same across the whole population, but rather dependent on the location of the sampled deme with respect to the origin of expansion. This is because the effect of colonization on intra-deme coalescence times becomes stronger with the distance from the origin of colonization (see Fig. 6, 7 in Austerlitz et al., 1997). If we assume that colonization has no strong effect on *S*_*ij*_, and considering that 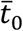 decreases towards the direction of expansion, then from the expression of *F*_*st*(*i,j*)_ (*Eq. 3*) we note that for *i* ≠ 0 and 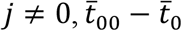 is greater than zero, increasing with the distance from the origin of colonization. Conversely, in an equivalent equilibrium model, 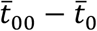 is always equal to zero. This might explain why, before HL&F, the range expansion simulation results show a larger effect of geographical distance on pairwise *R*_*st*_ (or *F*_*st*_) than an equilibrium model (Supplementary Appendix S1: Figure S11a, b). We can then conclude that in the range expansion scenario the influence of geography on pairwise *F*_*st*(*i,j*)_ is given by i) the relative contribution of *S*_*ij*_ on *F*_*st*(*i,j*)_, and ii) the spatial gradient in 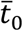 towards the direction of expansion, which, for demes far apart, results into pairwise *F*_*st*(*i,j*)_ that are higher than expected under an equilibrium model.

After full colonization of the habitat (before HL&F), 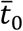 within each deme tends to the same equilibrium value over time, which is determined by the total population size of the continuous habitat (Austerlitz et al., 1997). Similarly, 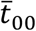 approaches to the value of a population at mutation-dispersal-drift equilibrium, such that 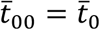. This means that differences in *F*_*st*(*i,j*)_ between equilibrium and range expansions scenario decrease significantly with the time since the onset of the expansion.

After HL&F, deme 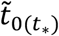 within each habitat patch tends to the same equilibrium value, given by the total population size of the habitat patch, which in our simulations is the same across patches. In the range expansion scenario, IBD is then expected to be lost i) once 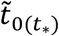 has reached the same value across habitat patches, regardless of their positions on the expansion gradient, and ii) once the relative contribution of the *spatial term* is much smaller than the *drift-isolation term*, as for the *instantaneous HL&F* scenario. Therefore, differences in *T*_*IBD*_ between *range expansion + instantaneous HL&F* and *instantaneous HL&F* scenarios could be explained by the fact that *T*_*IBD*_ in the *instantaneous HL&F* scenario only depends on the time required for the *spatial term* to become negligible compared to the *drift-isolation term*.

### Two-steps HL&F

It is probably more realistic to consider that habitat loss and fragmentation is a gradual spatial-temporal process. Therefore, in the present study we also compared the emerging IBD patterns in the case of an *instantaneous HL&F versus* a *Two-steps HL&F*. The scenario that we simulated consider a hierarchical HL&F process, where a first HL&F event generates three clusters of forest fragments which then undergoes a second HL&F event (Fig. 2b). The variable of interest in this case is the amount of time spent in the *intermediate state* between the first and the second HL&F events (*T*_*split*_). Our simulation results have shown that *Two-steps HL&F* can significantly influence the temporal dynamic of IBD patterns (Fig. 2b, Supplementary Appendix S1: Figure S8b, d and S10). The effect of *Two-steps HL&F* on IBD becomes stronger with increasing *T*_*split*_, since it takes more time for IBD to be lost for higher *T*_*split*_. Overall, this can be explained by the longer-in-time relative contribution of geography (*S*_*ij*_) on pairwise *R*_*st*_, the shorter *t*_*_ for habitat patches that were in the same cluster before the second HL&F event and by the slower decay in average coalescence time 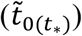. Note that, after 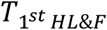, both connected (within the same cluster) and fragmented (different clusters) deme pairs show an increase in pairwise *R*_*st*_. However, pairwise *R*_*st*_ between demes that are within the same cluster after 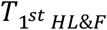 does not increase as much as demes that are at the same geographical distance but on different clusters (Supplementary Appendix S1: Figure S11c, d). It is therefore useful to first distinguish the processes that influence pairwise *F*_*st*_ for demes within or between clusters.

Increase in *R*_*st*_ for deme pairs within cluster can be deduced from the expression of *F*_*st*_ in a continuous habitat (*Eq. 2*), which can be re-written as 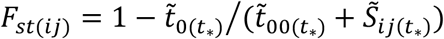 for a non-equilibrium population. Since the considered demes lie on the same cluster, 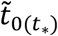 is equal to 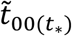, and decrease at the same rate after 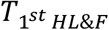. Also 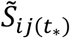 decreases after 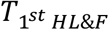, given that the amount of time required for two genes to be in the same deme (*S*_*ij*_; *Eq. 4*) is positively correlated to the total number of demes in the habitat. According to this *rationale*, an increase in *F*_*st*_ could then be explained by a slower decrease in 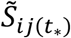 compared to 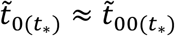. Increase in *R*_*st*_ for deme pairs between clusters can be explained following the arguments given for the *instantaneous HL&F*. In fact, under this condition, pairwise *F*_*st*_ evolve according to *Eq. 10*. However, we show that pairwise *F*_*st*_ between clusters increases slower than an equivalent comparison in the *instantaneous HL&F* (Fig. S11c, d). This is mainly due to the slower decrease in 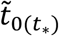 in the *Two-steps HL&F*, given that cluster size is about twice larger than patch size in the *instantaneous HL&F*.

Compared to an equivalent *instantaneous HL&F* scenario, the increase in *T*_*IBD*_ with *T*_*split*_ can thus be explained i) by the slower increase in pairwise *R*_*st*_ for both connected and fragmented subpopulations and ii) by the steeper IBD slope that is generated by the lower pairwise *R*_*st*_ of geographically close subpopulations, given that they were connected until 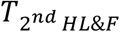

## Conclusions

The present study aimed at clarifying some aspects on the influence of HL&F on spatial genetic structure, taking into account both space and time. To our knowledge, few studies have theoretically addressed this question with spatially-explicit simulations (Ezard and Travis, 2006; Cushman et al., 2012; Mona et al., 2014; Jackson and Fahrig, 2016). More importantly, currently there are no analytical models in population genetics which jointly model space and time in the context of HL&F.

A study that is often cited is Hutchison and Templeton, (1999), where the authors proposed four hypothetical relationships between *F*_*st*_ and geographical distance, which had become, in many instances in literature, a reference for interpreting isolation-by-distance pattern from natural populations. Hutchison and Templeton, (1999) proposed four ‘verbal’ models for which they found support by analyzing genetic data from eastern collared lizards collected in four geographical regions in USA. This study has largely contributed to qualitatively classify the possible interactions between drift and gene-flow in species with geographically limited dispersal; however, it lacked a quantitative framework on which these hypotheses could rely on and being tested. With our study, we aimed to contribute and extend previous knowledge on the behavior of IBD in time and space, and provide an analytical framework to be explored according to the study case (e.g., landscape configuration).

Two of the theoretical study we cited (Cushman et al., 2012; Jackson and Fahrig, 2016) have investigated the effect of habitat amount or fragmentation *per se* (called ‘configuration’) on genetic structure through spatially-explicit simulations. The questions addressed in both studies aimed at identify if either habitat amount or habitat fragmentation *per se* would be better predictor of genetic structure. To do so, they performed individual-based simulations on landscapes with various degree of habitat amount and fragmentation *per se*, and correlated genetic structure (Mantel *r* values) with a range of fragmentation metrics. The two studies reached different conclusions. Cushman et al., (2012) found that habitat fragmentation has a higher effect on genetic structure than habitat area, whereas Jackson and Fahrig, (2016) found the opposite results. Jackson and Fahrig, (2016) discussed this discordance between the two studies considering, as also Cushman et al., (2012) point out in their work, that the fragmentation metrics used in Cushman et al., (2012) are strongly correlated with habitat amount. However, even Jackson and Fahrig, (2016) noted that although, overall, their results were pointing out to a minor role of fragmentation *per se* in shaping genetic structure, this was not correct for species with low dispersal rate. The analytical framework that we propose may help to such debate. Let’s first note that habitat fragmentation *per se*, defined as the breaking apart of a continuous landscape in habitat fragments, does implicitly reduce habitat amount. That is, once a complete barrier to gene-flow is formed (fragmentation) the only available habitat for a given population would be within the habitat fragment. The consequence is that the total population size of that population will be reduced 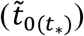, depending on the fragmentation configuration. Nevertheless, the total amount of habitat in the landscape do not necessarily change. In fact, the same reduction in total population size (and thus 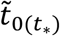 occurs even if that landscape is fragmented by sufficiently small barriers, such that landscape-scale habitat amount does not change. According to our model, an increase in pairwise *R*_*st*_ after habitat fragmentation is then mostly generated by the decay in average coalescence time within habitat fragments, suggesting that indeed total population size (within fragment) will play a major role in shaping genetic structure. However, this conclusion does not mean that habitat fragmentation has less importance than habitat amount, but rather that the two mechanisms are necessarily and often confounded.

## Limitations of the study and Future Directions

Our simulation results have shown that IBD patterns can persist for long time after HL&F. The demographic history of populations, prior to HL&F, can influence the time of IBD loss. This was shown with two examples: *range expansion before HL&F* and *spatio-temporal variation in HL&F*. In the present study, we did not fully explore the interaction between several possible parameters in the *range expansion* or *spatio-temporal variation in HL&F* scenarios, but rather provide a somehow proof-of-concept and intuition on how and to which extent the past of a given population can influence spatial genetic patterns. This question is not new (e.g., Nichols, 1996; Hewitt, 1999), but it was not yet explored in the context of HL&F (but see Mona et al., 2014). The results from more complex scenarios of HL&F have shown that, under certain conditions, an *instantaneous HL&F* model might not be appropriate. Our analytical model for *isolation-by-distance in time* assume the same mode of *instantaneous HL&F*. Still, the simplifying assumptions of the simulation and analytical model have helped providing a first and general understanding of how space and time jointly contribute to spatial genetic patterns. Moreover, it is not necessarily unrealistic to consider an *instantaneous HL&F*, if we consider the current rate and mode of HL&F in natural habitats due to anthropogenic activities (Silva Junior et al., 2020; Vancutsem et al., 2021). Nevertheless, our analytical model may result sufficiently accurate if used to interpret pairwise *F*_*st*_ between geographically close habitat fragments, since those are likely to behave as an instantaneous HL&F scenario.

The model could be further extended by considering for instance the work of (Austerlitz et al., 1997) on average coalescence time in a population under range expansion, and derive a framework that include both the impact of range expansion and HL&F on coalescence times and thus on pairwise *F*_*st*_. Other improvements could consist on overcoming the assumption of homogenous subpopulation size and dispersal rate across the landscape, or complete isolation among habitat fragments by considering permeable barriers to gene-flow (Ringbauer et al., 2018). In addition, the development of an inferential framework aiming at estimating time since HL&F (*t*_*_) based on, for instance, the method of moments or maximum-likelihood could be thought. Further extensions include also the use of the full distribution of coalescence times, instead of the expected coalescence times as done in the present study, which would allow then to explore more sophisticated information in the genome such as the distribution of genome block sharing (identity-by-descent blocks; IBD blocks) (e.g., Ringbauer et al., 2017).

## Author Contributions

GMS, LC and TM designed the project. GMS performed spatial genetic simulations and data analysis, and developed the numerical framework. TZ contributed to data analysis. TM and RR developed the genetic simulator. GMS drafted the manuscript and LC revised it.

## Acknowledgments

This work was supported by the 2015–2016 BiodivERsA COFUND call for research proposals, with the national funders Agence Nationale de la Recherche (grant number ANR-16-EBI3–0014), Fundação para a Ciência e Tecnologia (grant number Biodiversa/0003/2015) and German Bundesministerium für Bildung und Forschung (grant number 01LC1617A). This work was also supported by the Fundação para a Ciência e Tecnologia (grant numbers PTDC/BIA-BEC/100176/2008, PTDC/BIA-BIC/4476/2012, PTDC-BIA-EVL/30815/2017 to L.C., PD/BD/114343/2016 to G.M.S), by the Laboratoire d’Excellence (LABEX) entitled TULIP (grant numbers ANR-10-LABX-41 and ANR-11-IDEX-0002-02) as well as the the LIA BEEG-B (Laboratoire International Associé–Bioinformatics, Ecology, Evolution, Genomics and Behaviour) and the Investissement d’Avenir grant of the Agence Nationale de la Recherche (grant number CEBA: ANR-10-LABX-25-01). We thank Tiago Paixão and Montgomery Slatkin for useful discussions and comments on the projects.

