## Supplementary Appendix S1 for "The effect of habitat loss and fragmentation on isolation-by-distance and time"

### Supplementary Figures

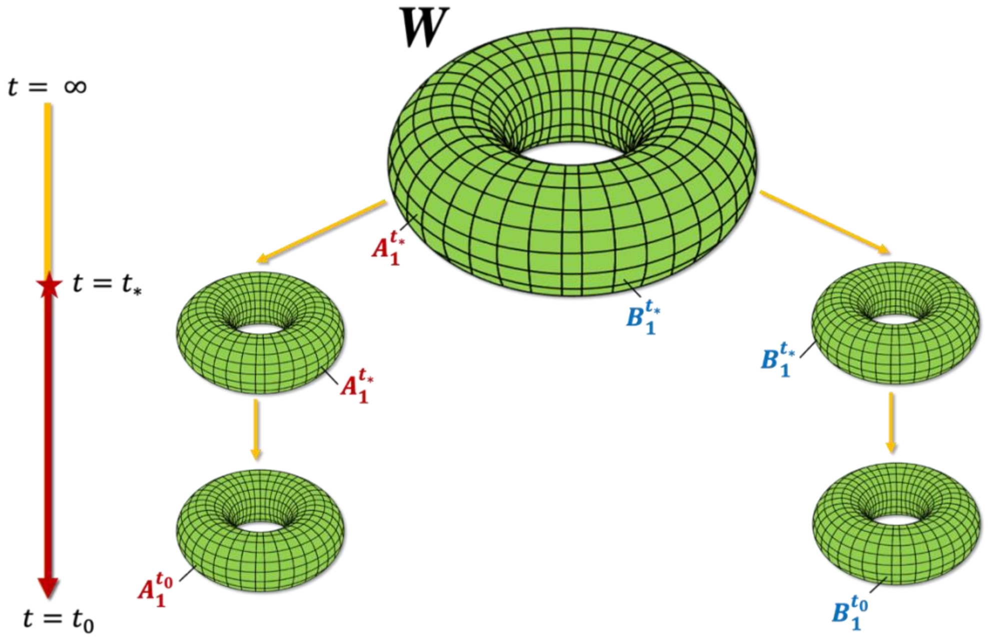


**Figure S1. Illustration of the isolation-by-distance in time model in the context of habitat loss and fragmentation.** $\boldsymbol{A}_1^{t_0}$and $\boldsymbol{B}_1^{t_0}$ indicate the location of individuals sampled in habitat fragments $A$ and $B$, respectively, at time $t = t_0$. $\boldsymbol{A}_1^{t_*}$ and $\boldsymbol{B}_1^{t_*}$ indicate one possible location of the ancestral lineages at the time of HL&F ($t = t_*$) for individuals sampled in habitat fragments $A$ and $B$, respectively. $W$: finite two-dimensional habitat prior to HL&F. $t = \infty$ specifies that population $W$ is at mutation-drift-dispersal equilibrium.

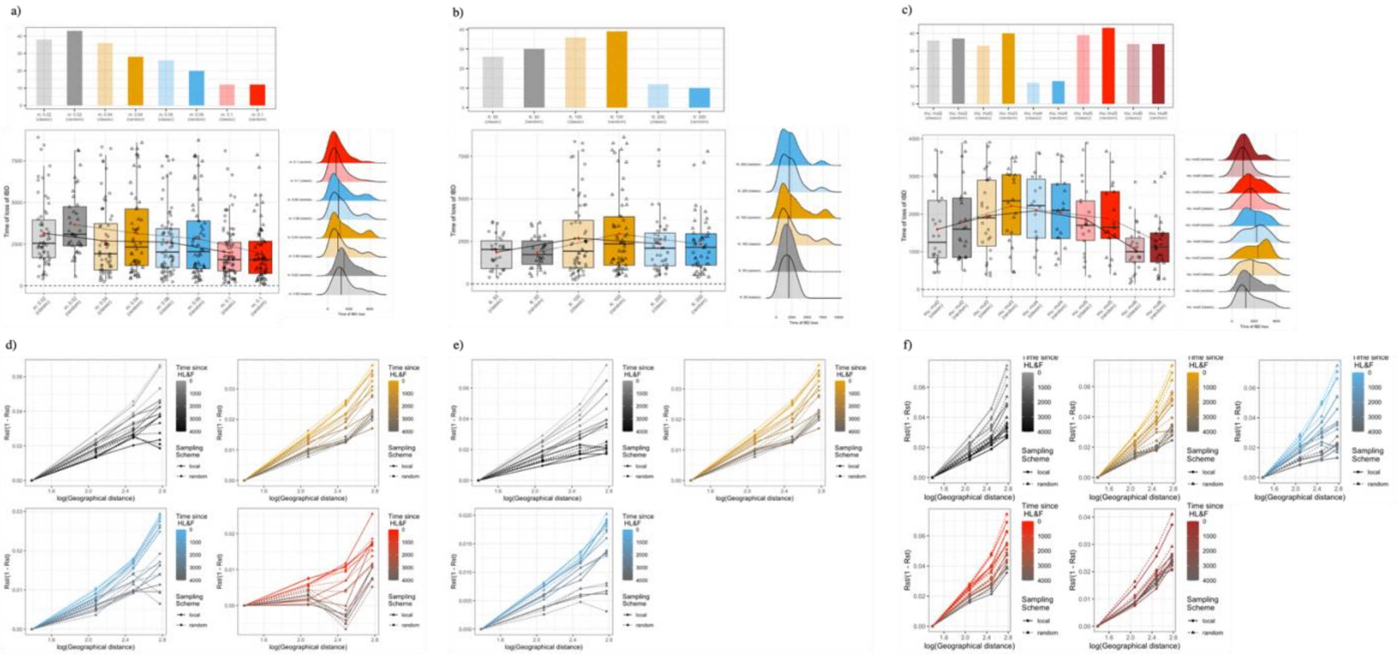


**Figure S2. The effect of HL&F on $R_{st}$-based isolation-by-distance (IBD) in relation to dispersal rate, deme population size and mutation rate.** Time at which IBD is no longer significant ($T_{IBD}$) as a function of **a)** dispersal rate ($m = 0.02$-$0.01$, $K = 100$, $\mu = 5 \times 10^{-4}$), **b)** deme population size ($K = 50$–$200$, $m = 0.04$, $\mu = 5 \times 10^{-4}$) and **c)** mutation rate ($\mu = 10^{-2}$–$10^{-6}$, $K = 50$, $m = 0.04$). Dots indicate the $T_{IBD}$ for each simulation replicate. Density plots are equivalent to box plot results. Upper histograms represent the number of replicates for which IBD was not lost within the time of the simulation. Mean pairwise $R_{st}$ as function of log(geographic distance) over time, distinguishing random (dashed line) and local (solid line) sampling in relation to **d)** dispersal rate, **e)** deme population size and **f)** mutation rate.

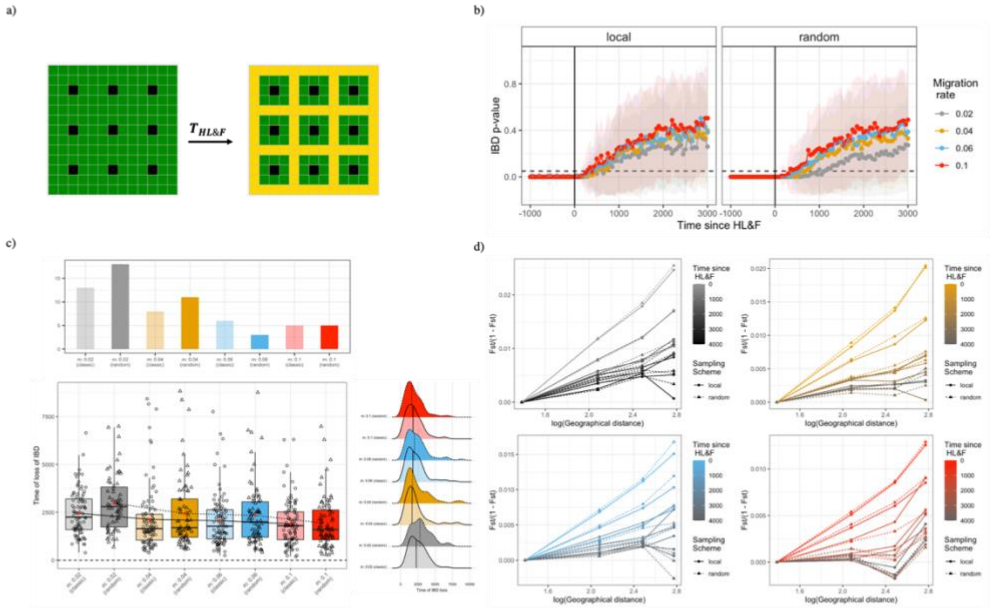


**Figure S3. The effect of habitat loss and fragmentation on $F_{st}$-based isolation-by-distance (IBD) in relation to dispersal rate. a)** Simulated 13 x 13 square grid of demes ($d_{tot} = 169$) undergoing HL&F at time $T_{HL\&F}$, sampling genetic data every 50 generations. Black cells refer to the sampled demes, green and yellow cells refer to suitable and unsuitable habitat, respectively. Dispersal rate values between 0.02 and 0.1 were tested, while fixing $K = 100$ and $\mu = 5 \text{ x } 10^{-4}$. **b)** Mean (dots) and standard deviation (shaded area) of IBD p-value across generations. **c)** Time at which IBD is no longer significant for each simulation replicate as a function of dispersal rate and sampling scheme (*local* vs *random*). Density plots equivalent to box plot results are shown for visualization purposes. Upper histograms represent the number of replicates for which no $T_{IBD}$ could be identified, because IBD was not lost within the time of the simulation. **d)** Mean pairwise $F_{st}$ as function of *log*(geographic distance) over time, distinguishing *random* (dashed line) and *local* (solid line) sampling. The results suggest that IBD is lost quicker at high dispersal rate.

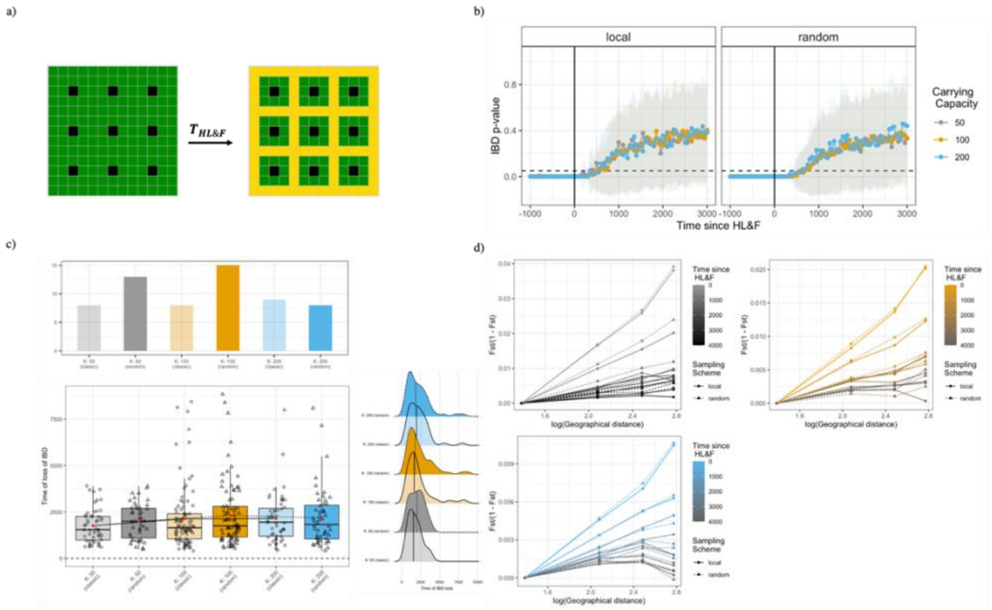


**Figure S4. The effect of habitat loss and fragmentation on $F_{st}$-based isolation-by-distance (IBD) in relation to deme population size. a)** Simulated 13 x 13 square grid of demes ($d_{tot}$ = 169) undergoing HL&F at time $T_{HL\&F}$, sampling genetic data every 50 generations. Black cells refer to the sampled demes, green and yellow cells refer to suitable and unsuitable habitat, respectively. Deme population size values between 50 and 200 were tested, while fixing $m$ = 0.02 and $\mu$ = 5 x $10^{-4}$. **b)** Mean (dots) and standard deviation (shaded area) of IBD p-value across generations. **c)** Time at which IBD is no longer significant for each simulation replicate as a function of deme population size rate and sampling scheme (local vs random). Density plots equivalent to box plot results are shown for visualization purposes. Upper histograms represent the number of replicates for which no $T_{IBD}$ could be identified, because IBD was not lost within the time of the simulation. **d)** Mean pairwise $F_{st}$ as function of log(geographic distance) over time, distinguishing random (dashed line) and local (solid line) sampling. The results suggest that deme population size has little effect on the time of IBD loss.

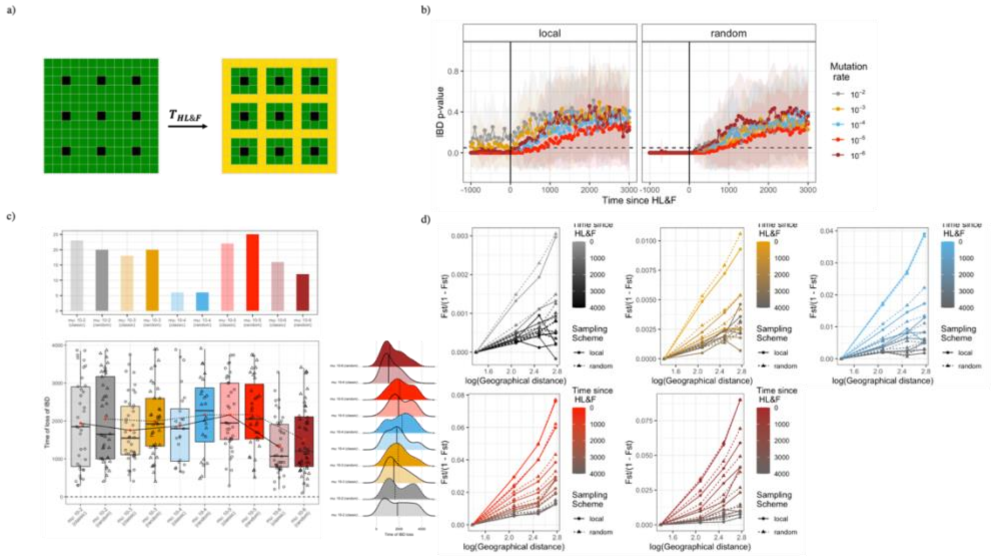

**Figure S5. The effect of habitat loss and fragmentation on $F_{st}$-based isolation-by-distance (IBD) in relation to mutation rate. a)** Simulated 13 x 13 square grid of demes ($d_{tot} = 169$) undergoing HL&F at time $T_{HL\&F}$, sampling genetic data every 50 generations. Black cells refer to the local sampling scheme, green and yellow cells refer to suitable and unsuitable habitat, respectively. Mutation rate values between 5 x $10^{-2}$ and 5 x $10^{-6}$ were tested, while fixing $K = 50$ and $m = 0.04$. **b)** Mean (dots) and standard deviation (shaded area) of IBD p-value across generations. **c)** Time at which IBD is no longer significant as a function of mutation rate and sampling scheme (local vs random). Density plots equivalent to box plot results are shown for visualization purposes. Upper histograms represent the number of replicates for which no $T_{IBD}$ could be identified, because IBD was not lost within the time of the simulation. **d)** Mean pairwise $F_{st}$ as function of log(geographic distance) over time, distinguishing random (dashed line) and local (solid line) sampling. The results suggest that IBD is lost quicker at high dispersal rate.

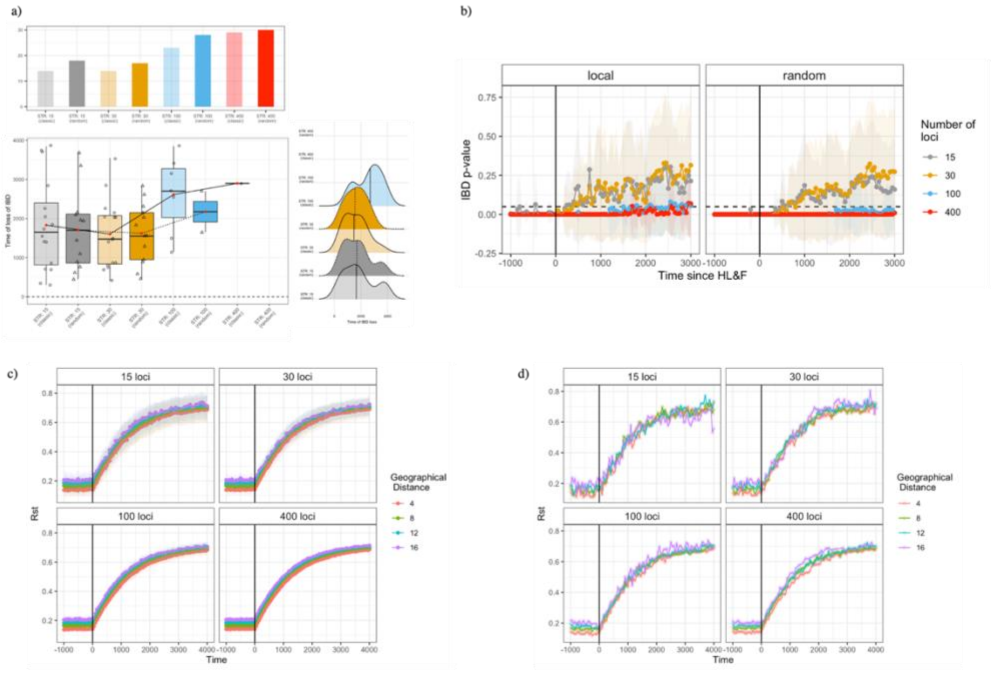


**Figure S6. Influence of the number of genotyped markers on the time of IBD loss since HL&F, based on $R_{st}$ isolation-by-distance. a)** Time at which IBD is no longer significant as a function of the number of markers (15 – 400) and sampling scheme (*local* vs *random*). Density plots equivalent to box plot results are shown for visualization purposes. Upper histograms represent the number of replicates for which no $T_{IBD}$ could be identified, because IBD was not lost within the time of the simulation. **b)** Mean (dots) and standard deviation (shaded area) of IBD p-value across generations. **c)** Pairwise $R_{st}$ over time as function of geographical distance averaged across simulation replicates (Mean: dots, standard deviation: shade). **d)** Pairwise $R_{st}$ over time as function of geographical distance for a single simulation replicate.

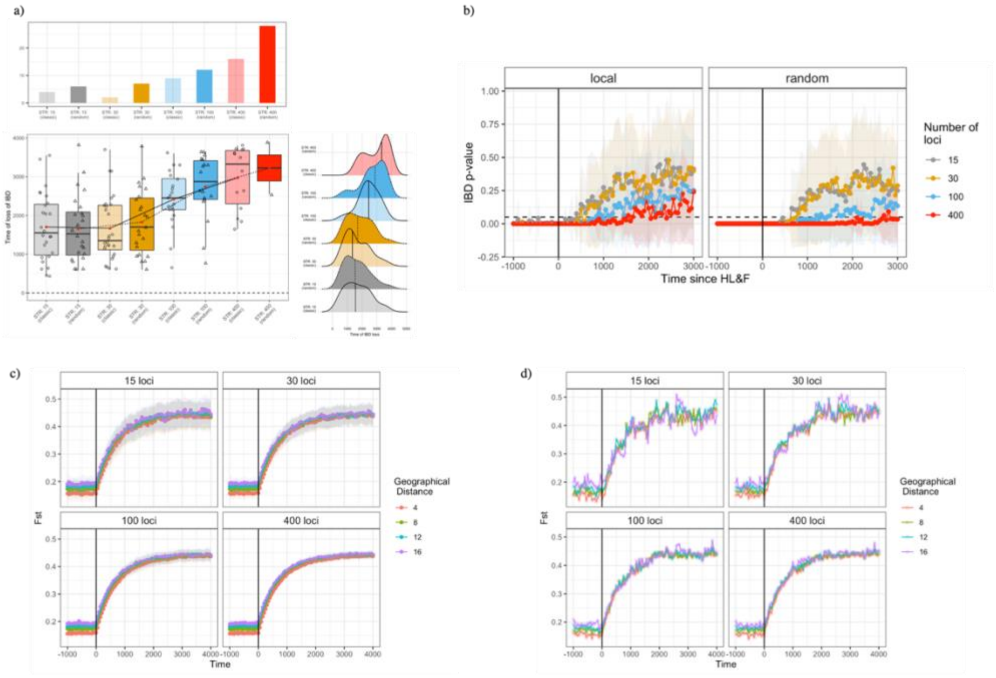


**Figure S7. Influence of the number of genotyped markers on the time of IBD loss since HL&F, based on $F_{st}$ isolation-by-distance. a)** Time at which IBD is no longer significant as a function of the number of markers (15 – 400) and sampling scheme (*local* vs *random*). Density plots equivalent to box plot results are shown for visualization purposes. Upper histograms represent the number of replicates for which no $T_{IBD}$ could be identified, because IBD was not lost within the time of the simulation. **b)** Mean (dots) and standard deviation (shaded area) of IBD p-value across generations. **c)** Pairwise $F_{st}$ over time as function of geographical distance averaged across simulation replicates (Mean: dots, standard deviation: shade). **d)** Pairwise $F_{st}$ over time as function of geographical distance for a single simulation replicate.

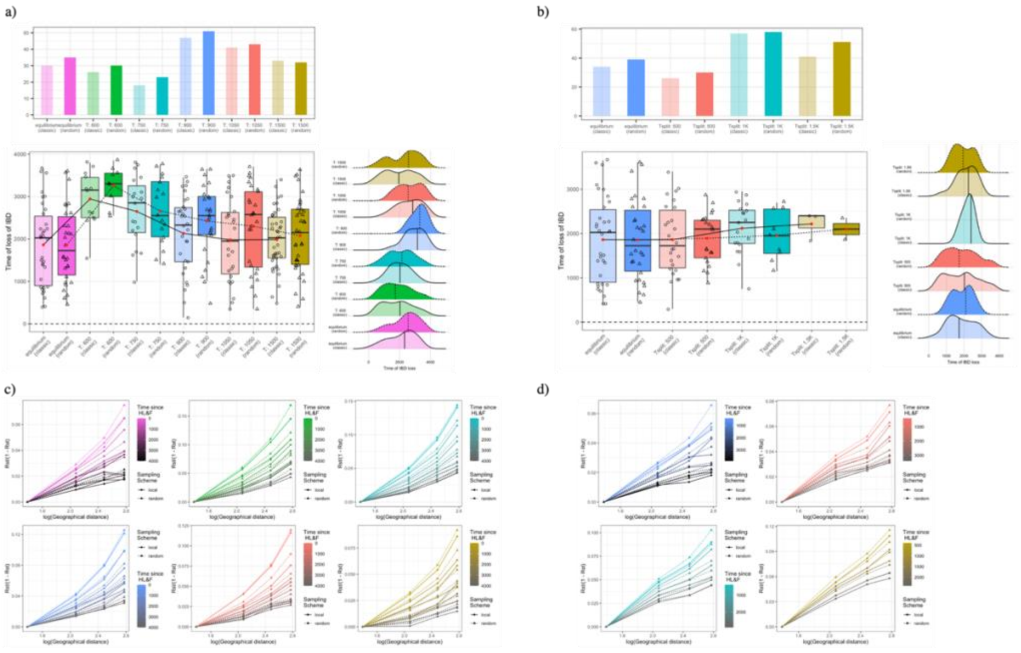


**Figure S8. The effect of ancient range expansion prior to HL&F and spatio-temporal variation in HL&F on loss of $R_{st}$-based isolation-by-distance (IBD).** Time at which IBD is no longer significant ($T_{IBD}$) for each simulation replicate as a function of **a)** time since the onset of the range expansion ($T_{exp}$ =60-1500, $K = 50, m = 0.04$ and $\mu$ =5 x $10^{-4}$), **b)** the time since the 1st HL&F event ($T_{split}$ =500-1500, $K = 50, m = 0.04$ and $\mu$ =5x$10^{-4}$). Density plots are equivalent to box plot results. Upper histograms represent the number of replicates for which IBD was not lost within the time of the simulation. Mean pairwise $R_{st}$ as function of log(geographic distance) over time, distinguishing *random* (dashed line) and *local* (solid line) sampling, and **c)** time since the onset of the range expansion or **d)** time since the 1st HL&F event.

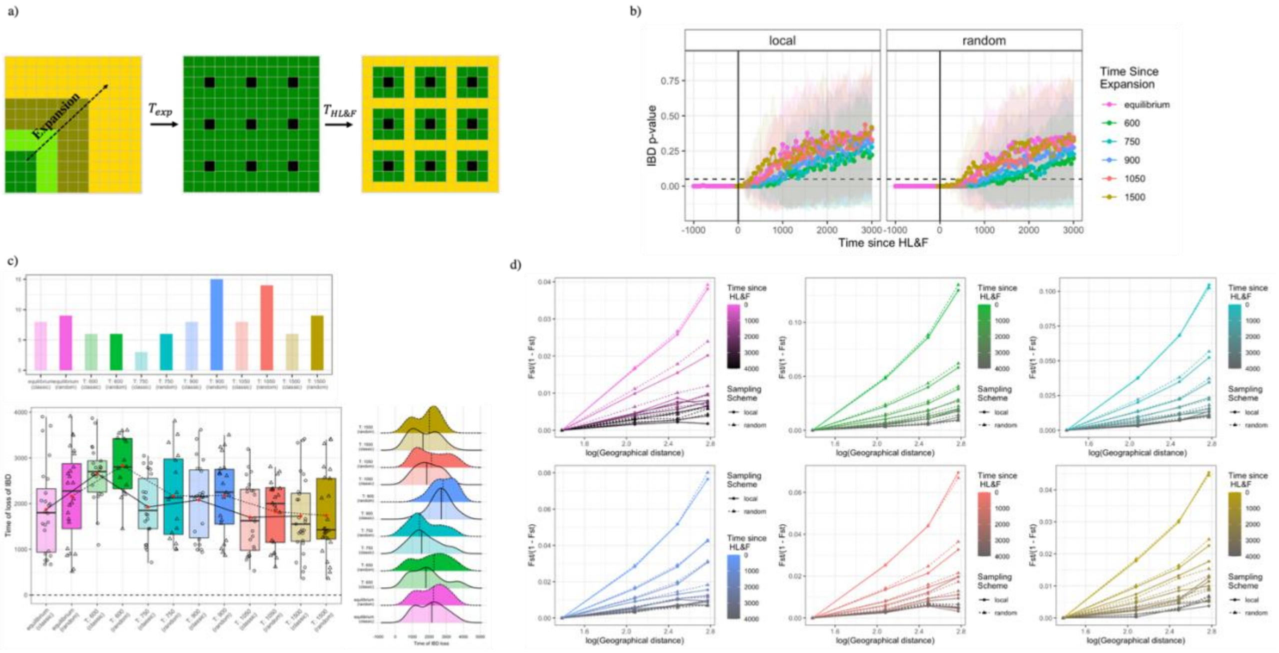


**Figure S9. The effect of ancient range expansion prior to HL&F on loss of $F_{st}$-based isolation-by-distance. a)** Simulated 13 x 13 square grid of demes, where $T_{exp}$ is the time since the onset of the range expansion and $T_{HL\&F}$ is the time of instantaneous HL&F. Range expansion was simulated for a scenario with $K = 50$, $m = 0.04$ and $\mu = 5 \times 10^{-4}$. **b**) Mean (dots) and standard deviation (shaded area) of IBD p-value across generations. **c**) Time at which IBD is no longer significant ($T_{IBD}$) for each simulation replicate as a function of sampling scheme (*local* vs *random*) and time since the onset of the range expansion. Density plots are equivalent to box plot results. Upper histograms represent the number of replicates for which IBD was not lost within the time of the simulation. **d**) Mean pairwise $F_{st}$ as function of log(geographic distance) over time, distinguishing *random* (dashed line) and *local* (solid line) sampling, and time since the onset of the range expansion.

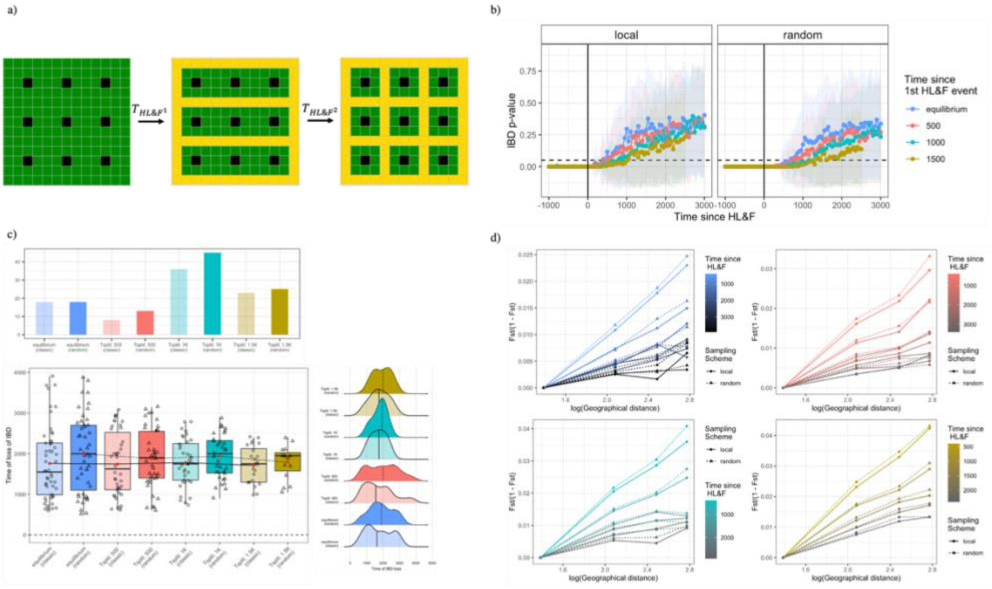


**Figure S10. The effect of spatio-temporal variation in HL&F on loss of $F_{st}$-based isolation-by-distance. a)** Simulated 13 x 13 square grid of demes, where $T_{HL\&F^1}$ and $T_{HL\&F^2}$ indicate, respectively, the time of the 1st and 2nd HL&F event. Piecewise HL&F was simulated for a scenario with $K = 50$, $m = 0.04$ and $\mu = 5 \times 10^{-4}$. **b**) Mean (dots) and standard deviation (shaded area) of IBD p-value across generations. **c**) Time at which IBD is no longer significant ($T_{IBD}$) for each simulation replicate as a function of sampling scheme (*local* vs *random*) and time since the 1st HL&F event. Density plots equivalent to box plot results are shown for visualization purposes. Upper histograms represent the number of replicates for which no $T_{IBD}$ could be identified, because IBD was not lost within the time of the simulation. **d**) Mean pairwise $F_{st}$ as function of log(geographic distance) over time, distinguishing *random* (dashed line) and *local* (solid line) sampling, and time since the 1st HL&F event.

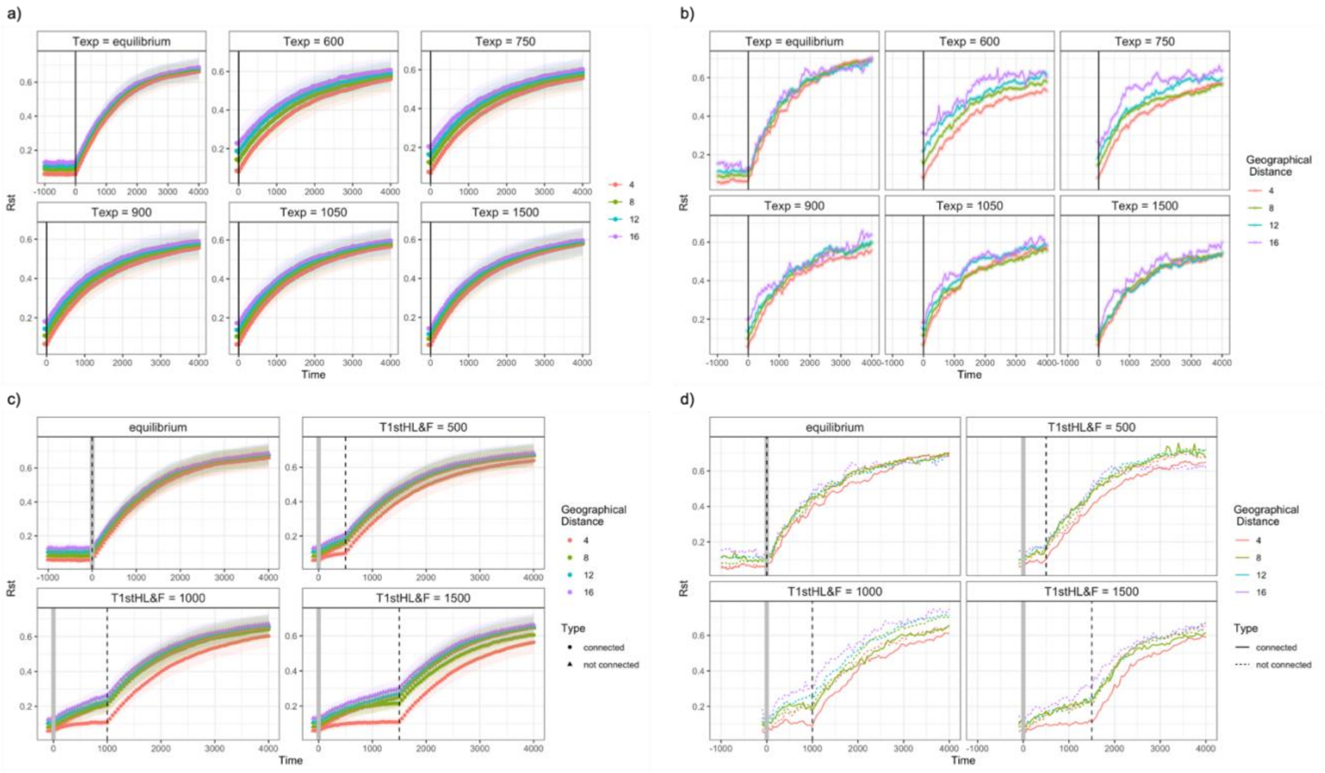


**Figure S11. The effect of range expansion prior to HL&F and spatio-temporal variation in HL&F on pairwise $R_{st}$.** First row: effect of the time since the onset of the onset of the range expansion ($T_{exp}$); Second row: effect of the time since the 1st HL&F event. In both scenarios, carrying capacity, dispersal rate and mutation rate were fixed at $K = 50$, $m = 0.04$ and $\mu$=5 x $10^{-4}$. Panels **a)** and **c)** represent the mean (dots) and standard deviation (shade) of pairwise $R_{st}$ across simulation replicates. Panels **b)** and **d)** indicate pairwise $R_{st}$ over time for a single simulation replicate.

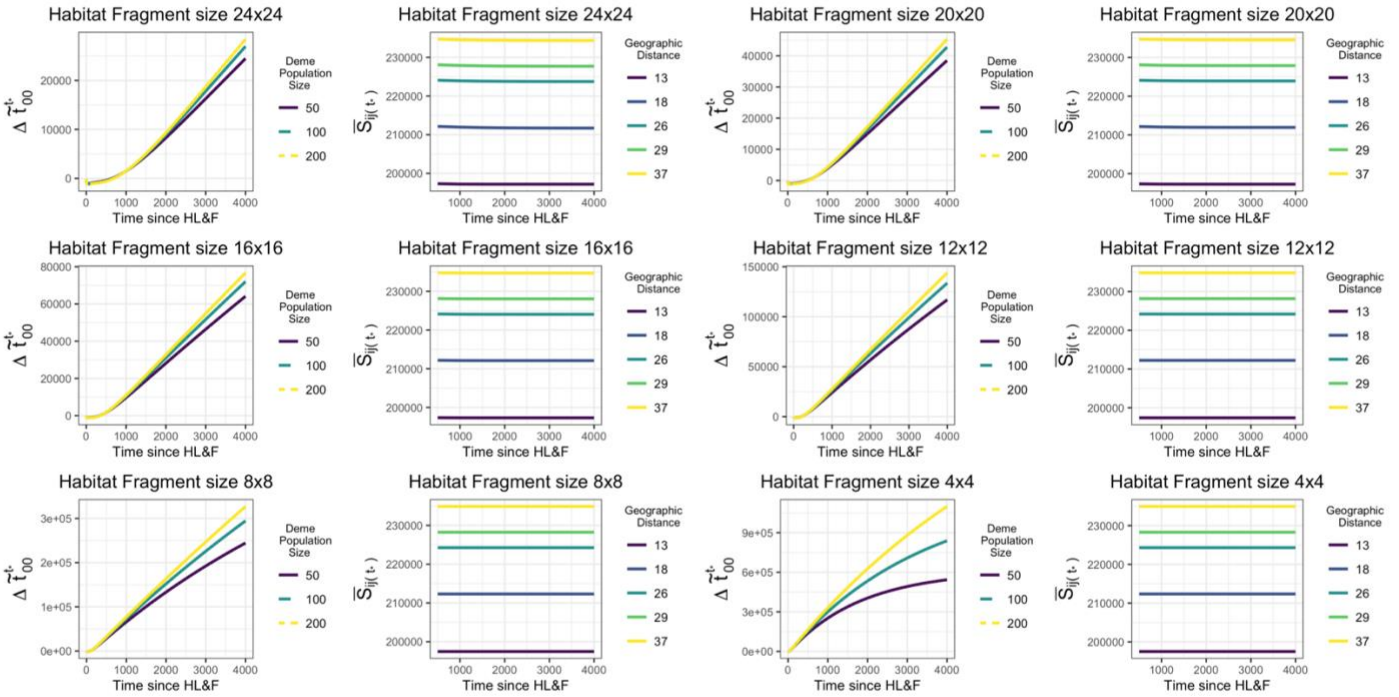


**Figure S12. *Spatial* and *drift-isolation* term dynamics after HL&F as function of deme population size and habitat loss.** An initial toroidal stepping stone model of size 78x78 is subject to HL&F, leading to the formation of nine torus-like fragments of size 24x24, 20x20, 16x16, 12x12, 8x8 or 4x4. Dispersal rate was fixed at $m = 0.02$. The *drift-isolation* term is given by $\Delta\tilde{t}_0^{t_*} \equiv \bar{t}_{00} + t_* - \tilde{t}_{0(t_*)}$ and the *spatial* term by $\bar{S}_{ij(t_*)}$. The results show that $\bar{S}_{ij(t_*)}$ remains nearly constant over time since HL&F and that $\Delta\tilde{t}_0^{t_*}$ changes are little affected by deme population size, expect for severe habitat loss (e.g., 4x4).

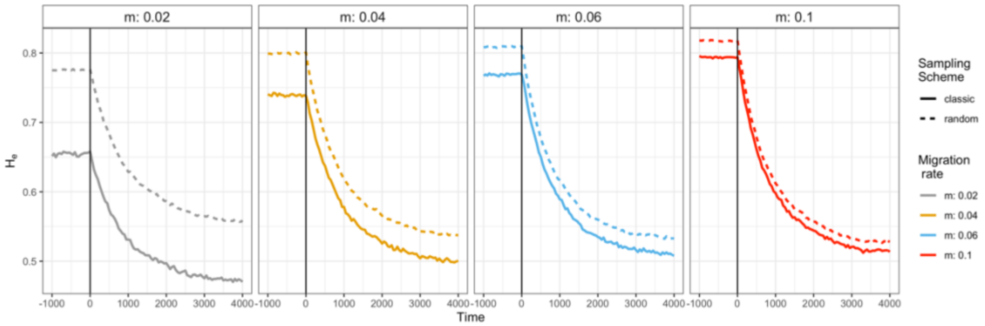


**Figure S13. Temporal dynamic of expected heterozygosity calculated using *random* or *local* sampling within habitat fragment.** Solid vertical lines indicate the HL&F event under the instantaneous HL&F scenario. Solid and dashed coloured lines refer, respectively, to *local* and *random* sampling. Across simulation, deme population size was fixed at $K = 50$. The results show that, at low dispersal rate, large differences in heterozygosity between sampling schemes can be identified, which tend to disappear with increasing dispersal rate.
